# A dynamical systems treatment of transcriptomic trajectories in hematopoiesis

**DOI:** 10.1101/2021.05.03.442465

**Authors:** Simon L. Freedman, Bingxian Xu, Sidhartha Goyal, Madhav Mani

## Abstract

Inspired by Waddington’s illustration of an epigenetic landscape, cell-fate transitions have been envisioned as bifurcating dynamical systems, wherein the dynamics of an exogenous signal couples to a cell’s enormously complex signaling and transcriptional machinery, eliciting a qualitative transition in the collective state of a cell – its fate. It remains unclear, however, whether the dynamical systems framework can go beyond a word-based caricature of the system and provide sharp quantitative insights that further our understanding of differentiation. Single-cell RNA sequencing (scRNA-seq), which measures the distributions of possible transcriptional states in large populations of differentiating cells, provides an alternate view, in which development is marked by the individual concentration variations of a myriad of genes. Here, starting from formal mathematical derivations, we challenge these transcriptomic trajectories to a rigorous statistical evaluation of whether they display signatures consistent with bifurcations. After pinpointing bifurcations along transcriptomic trajectories of the neutrophil branch of hematopoeitic differentiation we are able to further leverage the primitive features of a linear instability to identify the single-direction in gene expression space along which the bifurcation unfolds and identify possible gene contributors. This scheme identifies transcription factors long viewed to play a crucial role in the process of neutrophil differentiation in addition to identifying a host of other novel genetic players. Most broadly speaking, we provide evidence that, though very high-dimensional, a bifurcating dynamical systems formalism might be appropriate for the process of cellular differentiation and that it can be leveraged to provide insights. Ambitiously, our work attempts to take a step beyond data-analysis and towards the construction of falsifiable mathematical models that describe the dynamics of the entire transcriptome.

## 1. Introduction

During development and tissue regeneration, it is envisioned that cells progress through multiple transitions to ultimately adopt a distinguishable function. While each transition en-route to a terminal fate involves the coordination of myriads of molecules and complex gene regulatory networks interacting with external factors, there is a common view that they depend on significantly fewer control parameters. This view was notably explicated by Conrad Waddington, in an illustration of an epigenetic space as a tilted, bifurcating landscape, where a vast number of nodes (genes) provide the scaffold for the smooth hills and valleys (cell state) along which a pebble (cell) can reliably roll down until it finds a resting position (terminal fate) (Fig. 1A) (1).

**Fig. 1.**
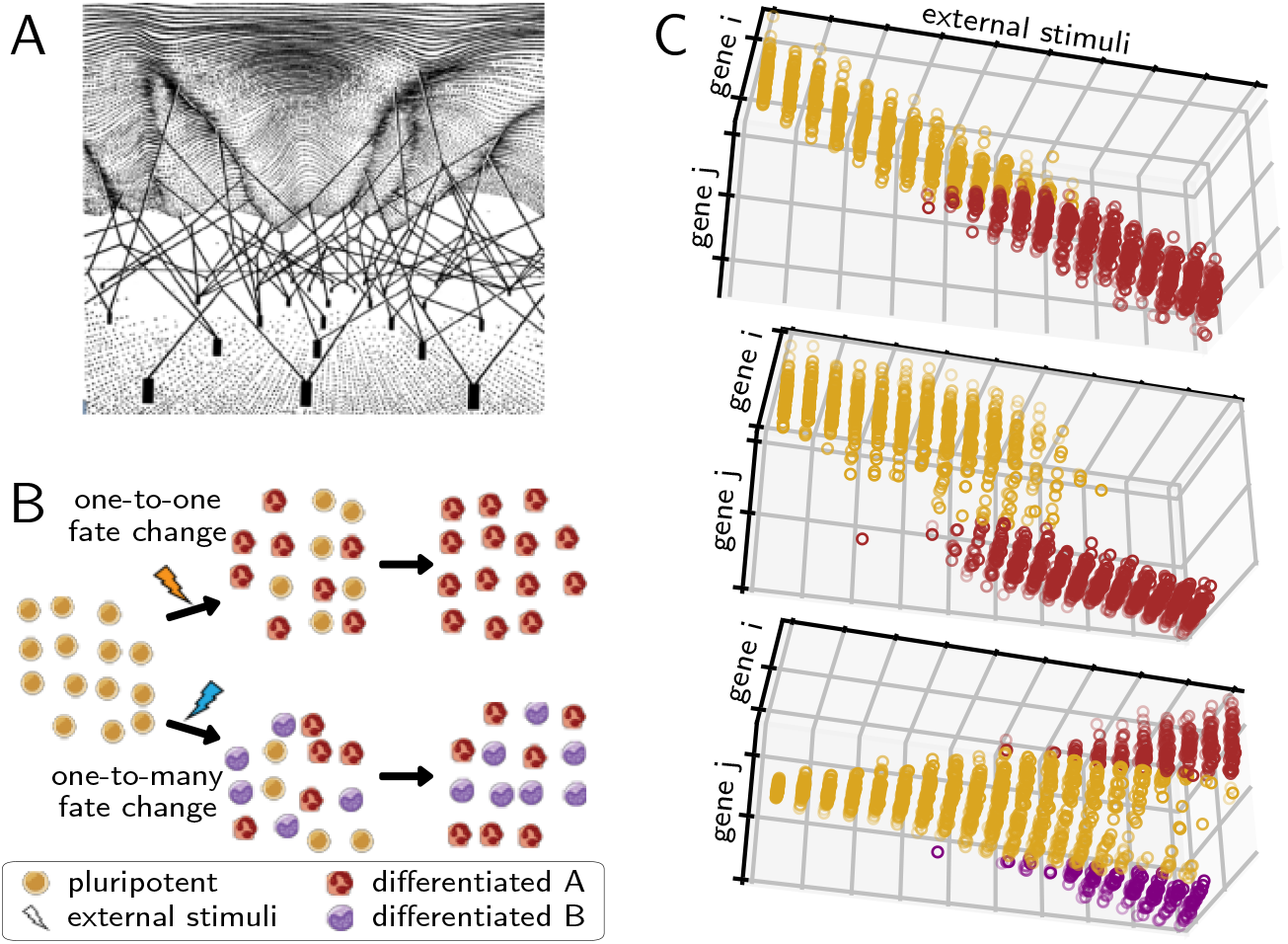
Cell type differentiation as a dynamical process. (A) Waddington’s landscape of cell fate commitment in which cell fates are represented as valleys, committment barriers as hills, and gene activity as pegs underneath that control the heights of hills and valleys. Adapted from Refs. (1, 17). (B) Schematic cell-population snapshots of maturation (top), in which one cell fate transitions to a different one, and a cell fate decision (bottom), in which a pluripotent cell differentiates to either of two lineages. Cell type images by Mikael Häggström, and A. Rad, used with permission and recolored. (C) Gene expression trajectories for cells (dots) at varying levels of a differentiating stimuli for cases where the differentiation landscape does not bifurcate (top), undergoes a saddle-node bifurcation (middle), or undergoes a pitchfork bifurcation (bottom).

Many of the characteristics of Waddington’s landscape have been codified into the language of dynamical systems, including that cell fates resemble valleys (attractors) in gene expression, or transciptomic, space (2–5), that a small amount of stable states can emerge from large interconnected boolean networks (6), and that known genetic interactions can yield multiple cell fates (bistability) (7, 8). Waddington’s illustration has motivated analysis of the wealth of data captured in single-cell RNA-sequencing (scRNA-seq) as well, in which the transcriptome of individual cells are measured, often at multiple time-points, as they differentiate. For example, a mathematical model of a pitchfork bifurcation can be fit to scRNA-seq data and yield predictions for developmental perturbations (9), large transcriptomic matrices can be dimensionally reduced to enhance the resolution of bifurcations to precisely determine the genes enabling a cell fate decision (10, 11), and well characterized cell-lineage relationships can be used to extract predictive models of gene regulation (12, 13). While these studies generally characterize cell fate decisions as bifurcations of an underlying developmental landscape, other studies model cell fate decisions as noise-induced state transitions on non-bifurcating developmental landscapes, by parameterizing cells’ dynamics and stochasticity, and use these models to infer lineage relationships and state transition probabilities (14–16).

In the face of these contrasting views, it remains unclear when, during development, transcriptomes undergo bifurcations. Even more abstractly, the strict geometric structure of bifurcations become less constraining as the dimensionality of the system grows, perhaps leading us to anticipate that the vast dimensionality of the transcriptome precludes the possibility of formal bifurcations being present, let alone found. However, the audacious hope of the perspective put forward in this study is that the same feature that permits ∼ 10^23^ atoms present in a straw to collude together to buckle at a critical load, the interactions amongst them, is present at a sufficient level within the transcriptome to display such collective phenomena. The central question guiding this study is therefore what statistical signatures must be present to confirm the presence of a dynamical system bifurcating and, once found, what additional insights does a bifurcation imply. Our goal is not to put forward a data-analysis tool, but instead to formally investigate the extent to which a bifurcating dynamical systems view of the transcriptome is an accurate and predictive mathematical description of the observed dynamics.

How is it possible to address these more abstract questions if single-cell RNA sequencing doesn’t usually involve a high-temporal-resolution measurement? To circumvent this short-coming of the measurement, and leveraging the inherent cellular variability in a population of differentiating cells, multiple groups have formulated pseudotime algorithms, parametric curve fitting tools that use the transcriptome similarity of cells to determine their relative ordering in developmental time, and thereby infer transcriptomic trajectories that span developmental decisions (10, 18, 19). The central assumption, which finds increasing empirical support (20– 23), is that the cell-level variability in a differentiating population is primarily along the path of cell-fate specification. Thus, stitching together the transcriptomes of a single population of cells assayed at the same lab time permits a dynamical measurement.

While recent work has used pseudotime as a proxy for developmental time, we take this further and show that it can be viewed as an axis along which a control, or bifurcation parameter(s) steadily varies, permitting a dynamical systems style investigation of possible bifurcations in the collective transcriptomic state. The dynamical systems view seems suitable due to the qualitative observation that the dynamic molecular processes that lead to changes in the transcriptomic state, such as signal transduction and transcription, generically occur on the order of seconds and minutes (24), while cell fates change over the course of hours or days (4), suggesting that developmental control parameters vary at significantly slower rates than their downstream molecular processes. This separation of timescales enables the possibility for cells to have steady transcriptomic states, and is therefore necessary for there to be bifurcations in development, since a bifurcation is a qualitative augmentation of the steady state solutions, or branches of a dynamical system.

Here, we layout and demonstrate a statistical formalism for detecting and interrogating bifurcations in developmental fate transitions directly from transcriptomic pseudotime trajectories. Contrasting previous studies (7–10), we do not assume any specific mathematical form for the underlying genetic interactions, nor do we assume the shape, or even existence of an underlying cell-fate landscape (Fig. 1B) (25, 26) since it is not our goal to discern a specific model. Instead we rigorously query whether the necessary statistical signatures of a bifurcations are present in a developmental timecourse. We build on and compare to similar styles of approach, which have been used to detect signatures and molecular mechanisms of disease (25, 27), analyze differentiation processes in temporal and pseudotemporal gene expression data (26, 28), and characterize reversibility in saddle-node bifurcations (29). We show that our data-driven approach enables us to distinguish between three different types of transcriptomic variation directly from systems-level data: a non-bifurcative cell fate change that is due to continuous changes in gene expression (Fig. 1C, top) without multistability; a cell fate change that is due to a one-to-one state transition (Fig. 1C, middle), such as those that may occur during terminal-fate maturation (30); and a cell fate change that is due to a one-to many-state transition like those that occur when pluripotent cells decide between multiple cell lineages (Fig. 1C, bottom). We apply our framework to a class of *in-silico* gene networks, to demonstrate its ability to recover the salient features of a bifurcating dynamical system, and examine the effects of high dimensionality and noise. We demonstrate the utility of our framework in the context of a recently published scRNA-seq exploration of hematopoiesis, and show that cell-fate bifurcations can be pinpointed and analyzed in scRNA-seq data even without detailed knowledge of the underlying system’s dynamics and controls. Finally, we demonstrate that our framework can be used to extract genetic relationships that are pivotal to the dynamics underlying a bifurcative cell-fate change, in order to recover genes of known importance in hematopoiesis and generate new testable hypotheses for genetic networks that underly a cell fate decision.

## 2. Theory: the relationship between the Jacobian and the Covariance

In this section we outline the theoretical foundations that enables our attempt to formally assess the presence and essential properties of bifurcations in high-dimensional transcriptomic dynamics. Summarizing our key insights, we layout a conceptual framework tailored to the analysis of transcriptomic data that, despite the absence of a mathematical model for the dynamics, can reveal and investigate the system’s most salient dynamical features, its bifurcations, from transcriptomic trajectories alone (26, 31).

A scRNA-seq measurement yields a transcriptomic matrix, where each row is a different cell and each column is a different gene (Fig. 2A). Various dimensionality reduction and visualization tools, such as t-SNE (32), and k-nearest neighbor maps (14), have been used to visualize this data and determine pairwise distances between cells based on their transcriptomes (for example, the methods described in Secn. S4.1). More recently, parametric curve fitting tools have been developed to determine how cells in transcriptomic space vary with respect to a state variable, such as developmental time (pseudotime) (10, 18, 19) (Secn. S4.1). Where a pseudotime trajectory contains a cellular state change, then one could perhaps determine which genes are responsible, by binning the cells along the trajectory to statistically compare **G**(*τ*), the transcriptomic matrix at pseudotime *τ*, with a neighboring matrix at a time Δ*τ* in the future, **G**(*τ* +Δ*τ*) (33). While such an analysis may reveal which genes exhibit dynamic signals, it will not reveal if the transcriptome’s dynamics have undergone a bifurcation, and may be sensitive to the discretization (13). Here, we present an alternative method, fundamentally rooted in dynamical systems theory and aimed at pinpointing cell-fate bifurcations, that only relies on being able to calculate **C**, the gene-gene covariance at a particular pseudotime (Fig. 2A).

**Fig. 2.**
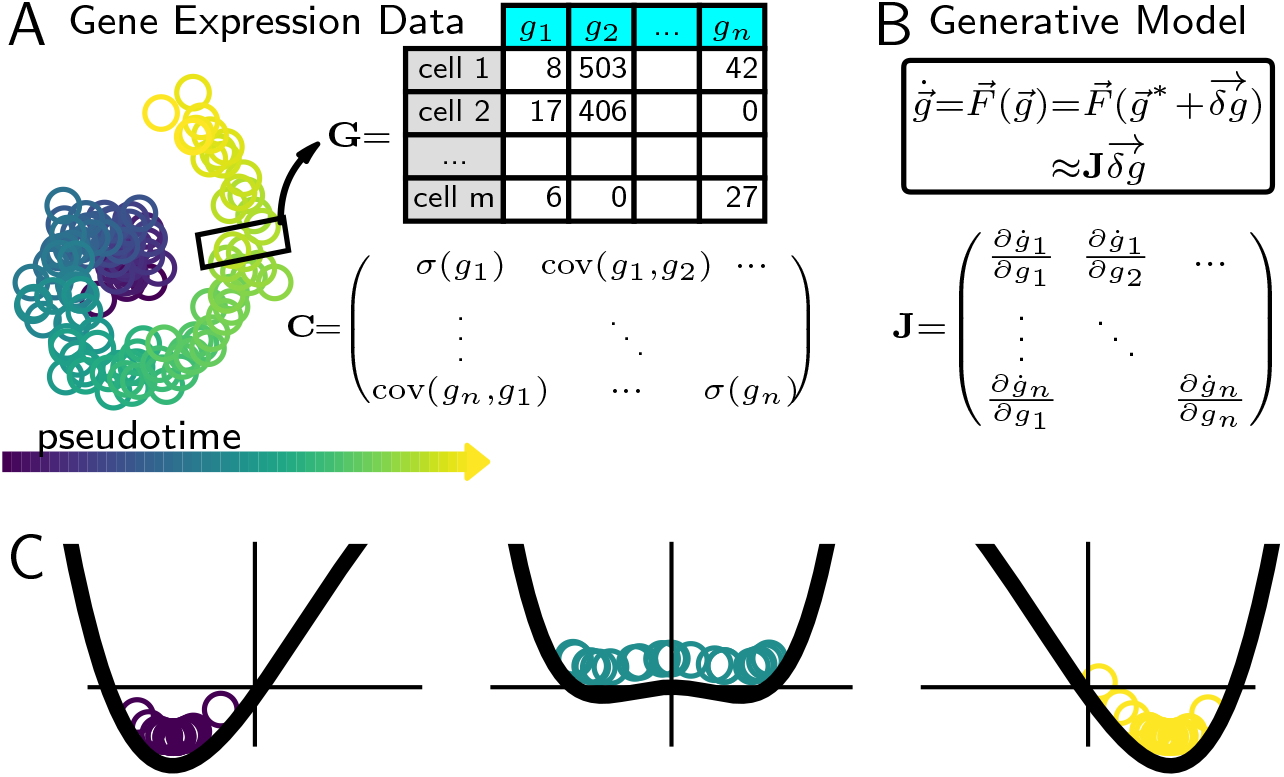
Relating the Jacobian to the Covariance at a bifurcation. (A) Schematic of a single-cell RNA-seq dataset arranged by each cell’s developmental (pseudo-) time. (Left) Visualization of dataset in two collective gene-expression dimensions. (Right) Gene expression matrix at the pseudotime indicated by the rectangle, and its corresponding covariance matrix. (B) Schematic of a generative model 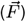 that could yield the gene expression matrix in (A), and its connection to the Jacobian (**J**). In this model, 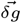 is the deviation of the gene expression vector, 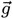 from the fixed point, 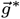. (C) Snapshots of a collection of particles at steady state following the dynamical process defined by 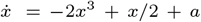 and uniformly sampled noise for *a* = − 1 (left), *a* = 0 (center) and *a* = 1 (right). Dark lines show the underlying landscapes 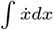.

Our analytical framework hinges on a fundamental mathematical relationship between the covariance matrix **C** estimated from a data matrix and the most salient properties of the underlying, unknown, generative model for the dynamics (illustrated in Fig. 2B). Regarding the underlying biochemical processes codified in a mechanistic model, we assume that (1) they are stochastic and Markovian (34–37),(15, 38) and (2) occur at significantly faster timescales (seconds to minutes) than the timescales over which transitions in cellular fates are observed (hours to days) (4, 24). A consequence of these assumptions (details in Secn. 5A) is that the local time evolution of a cell’s transcriptomic profile is controlled by a single matrix, the Jacobian (**J**), where *J*_*ij*_ = *∂ġ*_*i*_*/∂g*_*j*_ is the effect of the amount of gene *j* on the dynamics of gene *i* (Fig. 2B). Thus, the Jacobian matrix encodes the update rules that determine the future state of the transcriptome based on its current state and the exogenous signals presented to the cell. Note that the Jacobian matrix, in general, changes with pseudotime.

Under these assumptions, **J** relates to **C** through the continuous-time Lyapunov equation (39),

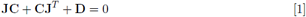

where **D** is the expected noise amplitude for individual genes and their interactions (derivation in Secn. 5A) (31). Eq. (1) is a remarkable relationship between the Jacobian and Covariance matrices of a system, implying that some mechanistic details of a system are encoded in, and recoverable from, the fluctuations of the state variables alone! While leveraging this relationship doesn’t make it possible to determine every element of **J**, since **D** is generically unknown and **J** (unlike **C**) is generically not a symmetric matrix, its most salient properties corresponding to its eigen-decomposition are inferrable from the covariance matrix near bifurcations.

Before we discuss the quantitative ramifications of this relationship in the context of a realistic high-dimensional dynamical system we first investigate a simple, but easy to visualize, one-dimensional toy-model, 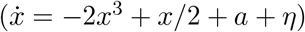 that undergoes a (imperfect pitchfork) bifurcation as a function of a control parameter (*a*) and is stochastic, controlled by the amplitude of the Langevin term, *η*. Fig. 2C displays the system before (*a <* 0), near (*a* = 0), and after (*a >* 0) the bifurcation. The slope of the potential function, drawn in black, determines the deterministic features of the dynamics of the system. In particular, parameter regimes (*a* ≪0 and *a* ≫0) where the potential has highly convex curvature exhibit stable fixed points, while parameter regimes near the bifurcation (*a* = 0), that have much flatter curvature exhibit instability. This dramatic reduction in curvature (mathematically, the curvature goes to 0 at the bifurcation itself) is the geometric signature of a bifurcation. Stochastic simulations of the system (drawn as open circles in Fig. 2C – color corresponds to the value of the control parameter) as a function of the control parameter demonstrate that owing to the reduction in curvature of the underlying potential, the data is spread maximally at the bifurcation, and narrows on either side of it.

The above toy model extends naturally to higher dimensions and captures the essence of the idea put forward in this paper. Summarizing the mathematical argument below, if a complex high-dimensional dynamical system undergoes a bifurcation, then in its vicinity there must be, by definition, some direction in the high-dimensional space whose curvature reduces dramatically, or softens. Consequentially, at the bifurcation point, the fluctuations in the system will be greatly enhanced along that soft direction. Generically, this direction need not point along any single dynamical variable (for example individual genes in a gene interaction network) of the original systems.

We now elaborate on the mathematical details of our central argument. Generically, a dynamical system’s local geometry can be obtained from diagonalizing **J**, such that,

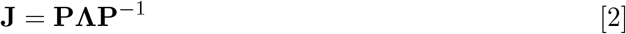

where Λ is a diagonal matrix of eigenvalues 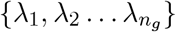 and **P**^*T*^ is the square matrix of eigenvectors 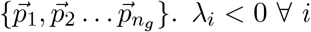 indicate a single, stable transcriptomic state, in the same way that the highly convex curvature in our toy model (Fig. 2C) indicate a single fixed point. Conversely, *λ*_*d*_ = max (Λ) → 0 indicates a bifurcation in the 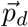 direction, just as the landscape of the toy model flattens out along the horizontal axis at *a* = 0 to enable the fixed point exchange. *λ*_*d*_ → 0 also considerably simplifies Eq. (1), such that it can be shown

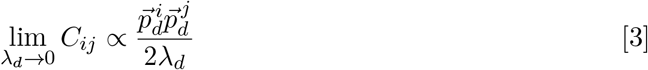

(details in Secn. 5B). The effect of *λ*_*d*_ → 0 on **C** is most clearly recognized by rewriting **C** in terms of its eigen-decomposition,

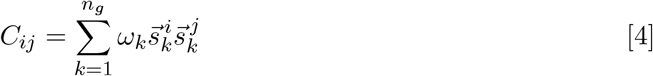

where 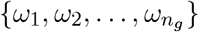 are its eigenvalues, and 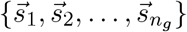 are its eigenvectors. Since all 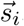, by definition, normalize to 1, Eq. (4) can only equal Eq. (3) if for at least one *ω* (*ω*_1_, without loss of generality)

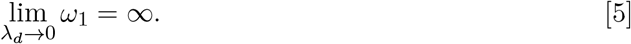

Thus, the covariance diverges along the principal direction 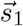, as illustrated for the toy model in Fig. 2C. Furthermore,just as the direction along which the variance expands in the toy model (*x*) is the same as the direction along which the landscape flattens, it can be shown, by equating Eq. (3) to Eq. (4) (where the *k* = 1 term will dominate; see Secn. 5C), that

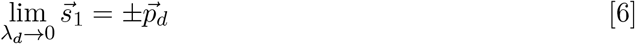

meaning, 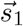 is the direction of the bifurcation! Lastly, we note a direct result of Eq. (3) is that at a bifurcation the Pearson’s correlation coefficient 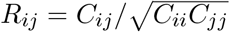 of the data along axes *i* and *j*, where their corresponding loadings on the eigen vector are non-zero, becomes maximal, as

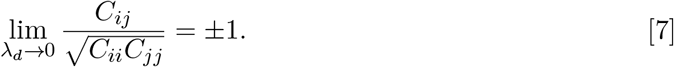

There is thus a secure correspondence between the fluctuations of a system and its dynamics. In particular, three specific changes to the transcriptomic covariance data, Eq. (5)-7, that can be determined from observations of the dynamics alone can inform us as to the salient features of the system, its bifurcations, even when we have no direct access to the generative model for the dynamics, or its corresponding underlying geometry. Notably, these features only rely on **G** being at steady state in the vicinity of an attractor, and do not rely on special circumstances, such as the Jacobian being symmetric, or the noise being of a particular nature. The expansion of correlation coefficients between crucial system features at a bifurcation (Eq. (7)) has previously been used to infer the networks that drive critical transitions (sometimes referred to as dynamic network biomarkers, or DNBs) (25, 27–29). More recently these bifurcation analysis methods have been used to identify critical developmental times from transcriptomic time-series (26, 40) and pseudotime trajectories (28). A notable difference in our analysis is our focus on the covariance eigen-decomposition, which directly reflects the dimensionality reduction that occurs at a bifurcation, rather than specific entries of the correlation matrix, which may require additional analyses to identify (see Secn. S1).

We first use theoretical models of noisy, high dimensional gene networks to concretely demonstrate how the approach outlined above (outlined in Secn. 7) can be leveraged to detect and assess bifurcations of an underlying dynamical system from observations of state-variables alone, bolstering previous applications. Following this, we emphasize the power of this approach by directly applying it to scRNA-seq data for the neutrophil lineage in the hematopoietic system.

## 3. Results

### A. Covariance analysis recovers salient features of a high-dimensional *in silico* gene regulatory network

To better understand our mathematical framework in the context of scRNA-seq data, where the large number of discordant genes and biological noise may obfuscate the predicted covariance signal indicative of a bifurcation, and cell fate changes may take different geometric forms, we tested the framework on a noisy, high-dimensional, gene-regulatory network (GRN), whose dynamics are governed by a set of explicit ordinary differential equations, **Ġ**= **F**(**G**), simulated with Poissonian noise (details in Secn. 6). In our model, cell fate transitions result from two mutually inhibiting “driver” genes, *g*_1_ and *g*_2_ via their dynamics

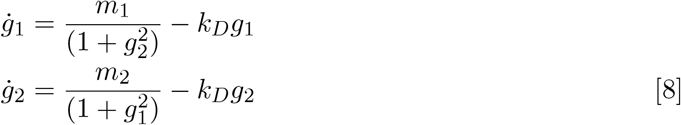

where *k*_*D*_ are their degradation rate, and *m*_1,2_ determine the scales of their synthesis (41). This simple system is illustrated in Fig. 3A. Varying the control parameter *m*_1_ yields a saddle-node bifurcation in gene-expression while varying *k*_*D*_ yields a pitchfork bifurcation (Fig. S1). Similar networks have been analyzed to provide insight into gene-inhibition and activation (42) in a diversity of biological systems, such as the lac-operon (43) and cell-cycle control (44).

**Fig. 3.**
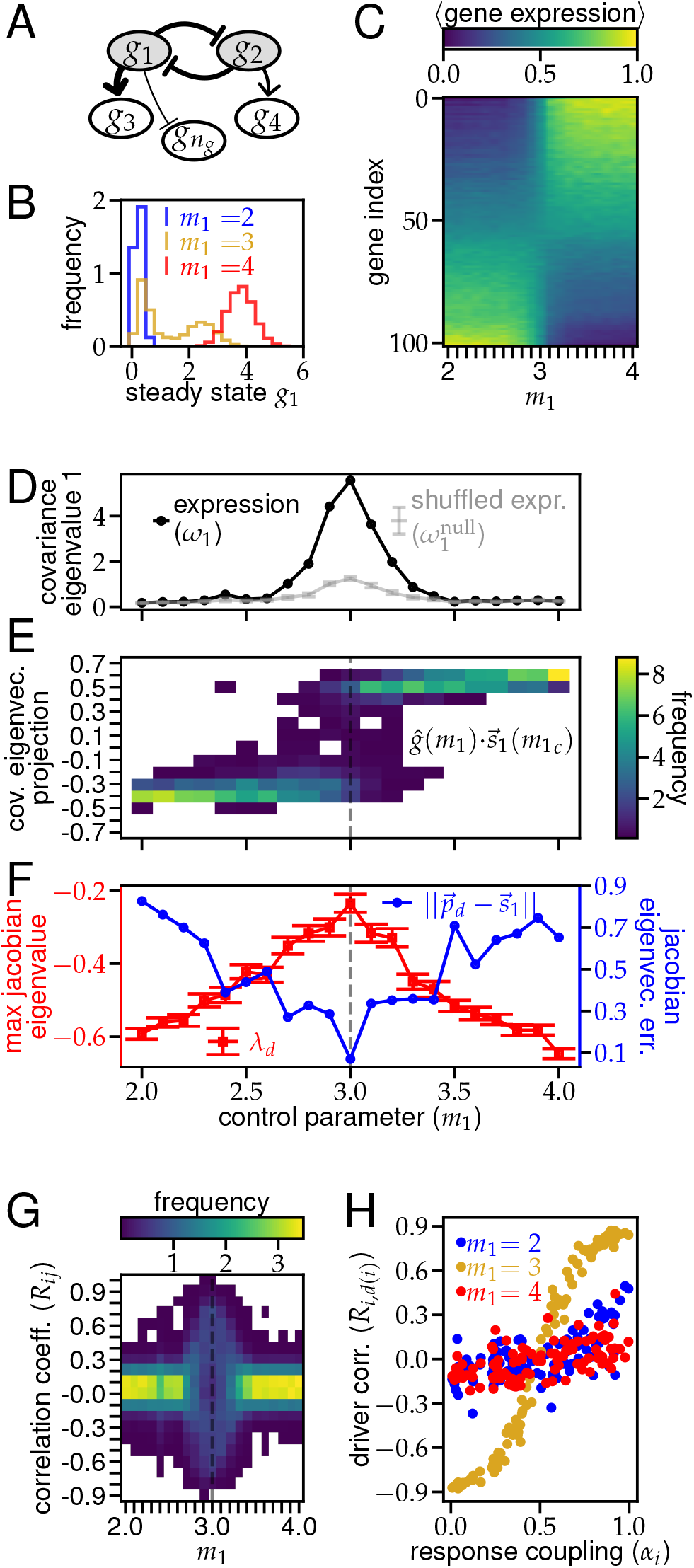
Analysis of a gene regulatory network around a saddle-node bifurcation. (A) Schematic of the GRN for 5 of the 102 genes. Undisplayed nodes have a unidirectional arrow stemming from either *g*_1_ or *g*_2_. (B) Distributions of *g*_1_ at steady state (see Secn. 6) for three values of *m*_1_. (C)-(G) GRN observables as a function of the bifurcating variable *m*_1_ evaluated over a distribution of 100 cells (Secn. 6). (C) Average final expression for each gene. Expression of driver genes, *g*_1,2_, (bottom and top rows, respectively) are min-max normalized. Response genes are sorted by their corresponding driver (*d*(*i*)) and activation level (*α*_*i*_). (D) Largest eigenvalue of covariance matrix shifted to have 0 min. (E) Distribution of normalized gene expression for each cell projected onto the bifurcating axis. (F) Red squares: largest eigenvalue of the Jacobian matrix. Error bars are SEM. Blue circles: Euclidean distance from the corresponding Jacobian eigenvector 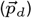 to the principle covariance eigenvector 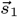. (G) Distribution of Pearson’s correlation coefficients for all gene pairs. (H) Correlation coefficients between responding genes and their driver as a function of the coupling coefficient *α* (Eq. (9)) at three values of *m*_1_. Each column in (E) and (G) integrates to 1.

As GRNs typically involve hundreds of genes, we include an additional *n*_*g*_ − 2 genes in the network that respond variably to the driver genes, according to

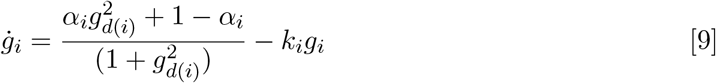

where *i* ∈ [3, *n*_*g*_]. *k*_*i*_ is the degradation rate of *i*^th^ gene, *g*_*i*_. The variable *d*(*i*) is set to 1 if *g*_*i*_ responds to *g*_1_ and 2 if it responds to *g*_2_. *α*_*i*_ ∈ [0, 1] is the strength of the connection between the responder *g*_*i*_ and its driver gene, *g*_*d*(*i*)_ (*α*_*i*_ = 0 yields full inhibition and *α*_*i*_ = 1 yields full activation). Thus each of these responder genes is connected to one of the two driver genes, to greater or lesser extents, as indicated in Fig. 3A. Though this GRN can be made more complex by including feedback from the responding genes to the two driver genes, or increasing the number of driver genes themselves, this simple model provides an interpretable demonstration of our proposed scheme.

We simulated Eq. (8)-Eq. (9) for a fixed number of genes (*n*_*g*_), statistical replicates, or cells, (*n*_*c*_), noise scale (*s*), duration (*N*_*t*_), and timestep (*δt*) for different values of the control parameters (*m*_1_, *k*_*D*_) (details in Secn. 6). We define the [*n*_*c*_ × *n*_*g*_] transcriptomic matrix **G**(*m*_1_, *k*_*D*_) once the system has reached steady-state in the simulation. We observed that the steady state distributions for individual genes (for example, *g*_1_, shown in Fig. 3B) shift their mean as the control parameter, *m*_1_, is varied, and exhibit bimodality at the bifurcation point, *m*_1_ = *m*_1*c*_ = 3, as expected for saddle-node bifurcations (Fig. 1B). We verified that *N*_*t*_ was sufficiently large by averaging **G**(*m*_1_) across cells, and observing that individual genes discontinuously, but predictably, switch their expression at *m*_1*c*_ (Fig. 3C; genes sorted by *d*(*g*_*i*_) and *α*_*i*_) compared to the continuous and unpredictable transitions observed with low *N*_*t*_ (Fig. S2A).

Having verified that our model simulates a system that undergoes a high dimensional saddle-node bifurcation driven by a 2-gene driver core, we use it to examine the effects of noise and a large number of responding genes on the theoretical predictions described in Secn. 2 (Eq. (5),6,7). As predicted, we found that *ω*_1_(*m*_1_), the largest eigenvalue of the covariance of **G**(*m*_1_), is maximal at the critical value *m*_1*c*_ (darker line in Fig. 3D), and the increase is significantly larger than can be obtained from a null distribution (lighter line in Fig. 3D). We generated this null through a marginal-resampling approach, which explicitly destroys any correlation between genes of **G**(*m*_1_) (details in Secn. S3). This contrast between the data and the null can be understood by considering the bimodality of the transcriptomic distribution at the bifurcation. Far from the bifurcation, the transcriptomic distribution is unimodal, and all *ω*_*i*_ scale with the noise scale *s*, which is undirected, and therefore unaffected by resampling, yielding 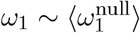 (Fig. S3 left and right panels). However, at the saddle-node bifurcation, the transcriptomic distribution is bimodal, so *ω*_1_ scales with the distance between the two modes (Fig. S3 center-top); marginal resampling of transcriptomes at the bifurcation yields new modes and the increased dimensionality of the bifurcation diminishes 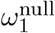, compared to *ω*_1_ (Fig. S3 center-bottom). While Fig. S3 only demonstrates the bifurcation bimodality in *g*_1,2_, the full transcriptomic bimodality can be visualized by computing 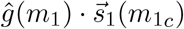, the normalized projection of each cell’s transcriptome along the principal covariance eigenvector. The distribution of this projection is densely centered around different fixed points to the right and left of *m*_1*c*_, but widens significantly at *m*_1*c*_ as there is non-zero probability for both transcriptomic modes (Fig. 3E).

Since, in this example, we have an explicit generative model (**F**(**Ġ**) given by Eq. (8)-9), we can validate that just as *m*_1*c*_ resembles a bifurcation from analysis of the covariance matrix, it also resembles a bifurcation of the full, noisy GRN, from analysis of the Jacobian. We show that the maximum, negative, eigenvalue (*λ*_*d*_) of the Jacobian 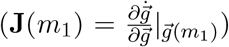 for this network approaches 0 from below as *m*_1_ → *m*_1*c*_ (Fig. 3F). We also show that at *m*_1*c*_, the direction of maximal covariance is given by the corresponding eigenvector of the Jacobian 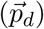, as the Euclidean distance between 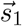, the principal eigenvector of the covariance, and 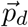 approaches 0 as *m*_*c*_ → *m*_1*c*_ (Fig. 3F). Thus, while the finite system size (*n*_*c*_) prevents, or regularizes, *ω*_1_ from diverging, and 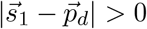, *ω*_1_ is still at its largest and the eigenvectors are in closest correspondence, at the bifurcation.

While the covariance eigen-decomposition provides insight into the timing and direction of a bifurcation, the theory predicts that the (Pearson) correlation coefficients between genes may help determine which genetic relationships are most critical for the dynamics at the bifurcation. We found that for low and high *m*_1_, when the network only has one fixed point, the distribution of *R*_*ij*_ is strongly centered around 0 (Fig. 3G). However, at the bifurcation, this distribution spreads out to ± 1 as predicted in Eq. (7) (Fig. 3G). To determine if the gene pairs that yielded large *R*_*ij*_ corresponded with critical gene relationships in our network, we plot *R*_*i,d*(*i*)_ the correlation between all responder genes and their drivers, sorted by their connection strength *α*_*i*_. We demonstrate that these correlation coefficients were much more strongly indicative of the responder-driver dependency (*α*_*i*_) at the bifurcation (Fig. 3H, green) than far away from the bifurcation (Fig. 3H, red and blue). Again, the model makes explicit that although the correspondence between geometry and dynamics is not universal, owing to the high-dimensionality of the system, in the vicinity of a bifurcation, a form of dimensionality reduction emerges which enables gleaning geometric characteristics directly from the dynamics. Thus, entries of a correlation matrix with high magnitude at a bifurcation may be reliable indicators of mechanistic gene-regulatory features.

We further used our model to understand Eq. (5) in the context of the pitchfork bifurcation induced by varying *k*_*D*_ (Fig. S4A). Unlike the example of a saddle-node bifurcation, we observed that *ω*_1_ does not peak at the bifurcation parameter *k*_*Dc*_ = 0.5, but rather begins to increase (Fig. S4B). This feature directly follows our interpretation that *ω*_1_ corresponds to the distance between the two modes of the transcriptomic distribution. While the bimodality of the saddle-node bifurcation results from the discontinuous transition between states, the bimodality of a pitchfork bifurcation emerges continuously from its root and becomes more pronounced as the control parameter is increased. Therefore, the distance between the modes (*ω*_1_) increases with the control parameter. By clustering the cells according to their transcriptomic mode, or branches, we are able to recover the bifurcation signature predicted by Eq. (5) (Fig. S4C), but we note that precise clustering requires prior knowledge (e.g., of how many clusters there are).

In this example, we have demonstrated the applicability and power of the theoretical infrastructure outlined above to analyze a high-dimensional and noisy dynamical system undergoing a variety of bifurcations, by uncovering its crucial aspects, including its location, direction in gene space, and influential genetic relationships. These calculations are also computationally simple; while covariance matrices can be cumbersome to compute for large numbers of genes and cells, reduced singular value decomposition can be used to quickly determine its largest eigenvalue and eigenvector, which is all our approach requires. Notably, our results only apply if the system is measured at steady state; in the absence of which there is no reason to anticipate clear divergences in the distribution of eigenvalues, transient bimodality, or equivalence between the covariance and Jacobian principal directions (Fig. S2B-D).

### B. Covariance analysis pinpoints a bifurcation in Hematopoietic stem cell development

Having verified that gene-gene covariance can be used to identify and classify a bifurcation in a simulated genetic context, we applied our analysis framework to a recently published scRNA-seq data set of mouse hematopoietic stem cell (HSC) differentiation (23).In this experiment, HSCs were isolated *in vitro*, barcoded, plated in a media that supports multilineage differentiation (day 0), and subsequently sampled for single-cell sequencing using inDrops (45, 46) on days 2, 4, and 6. The resultant transcriptomic matrix (25,289 genes in 130,887 individual cells) was filtered to only include highly variable, non-cell cycle genes, and visualized in 2*D* via the SPRING method (14) (Fig. 4(A)), in which the (*x, y*) coordinates of each cell are determined by optimally placing each cell closest to its 4 nearest neighbors in the space of the top 50 gene-wise principal components (PC) (14). Each cell was then ascribed to one of 11 different cell types (annotations in Fig. 4(A)) based on its position in the SPRING plot and expression of cell-type specific marker genes (23). Cells that belonged to the developmental transition from multipotent progenitor (MPP) to neutrophil were identified by recategorizing cells as a cell-label distribution, and ranking cells by their similarity to fully committed neutrophils (details in Secn. S4.1). The 61,310 cells identified as belonging to the neutrophil transition were sorted into a neutrophil pseudotime trajectory (Fig. 4A) by ranking cells according to their similarity with the earliest pluripotent cells (Secn. S4.1). This data-specific pseudotime algorithm was validated via the clonal barcodes, by ensuring that the MPP cells in the trajectory included neutrophil clones, and via the sequencing time, by ensuring that cells collected earlier were ranked earlier in the trajectory. We found that expression of hundreds of highly expressed and variable genes in these cells varied temporally along this trajectory, with large groups of cells either monotonically increasing or decreasing (Fig. 4B).

**Fig. 4.**
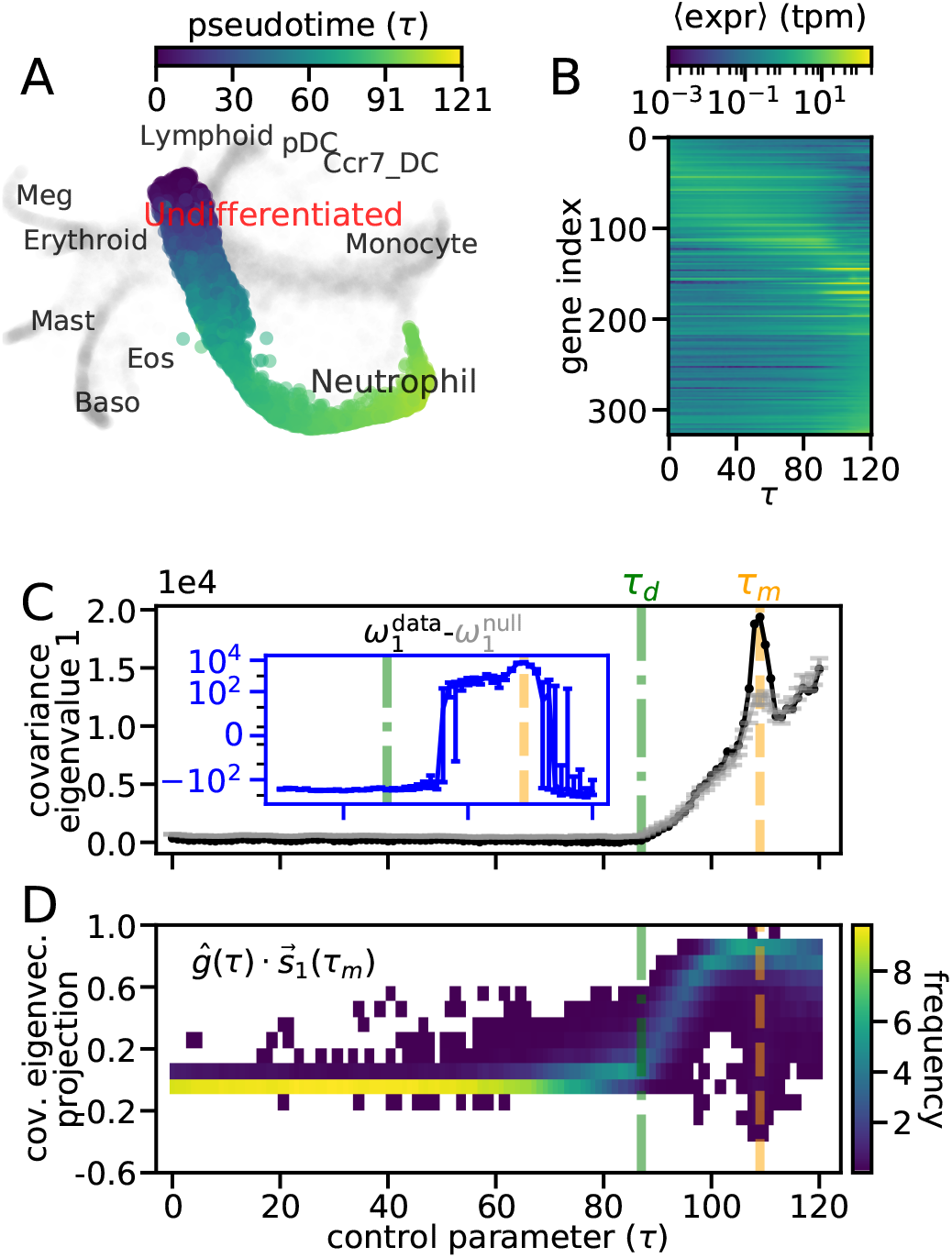
Covariance analysis of temporal scRNA-seq data shows signatures of developmental bifurcations. (A) SPRING visualization for each cell (point) in an *in-vitro* scRNA-seq experiment of mouse hematopoeitic stem cell differentiation (23). Cells in the pseudotime trajectory analyzed are colored accordingly (blue to yellow) while others are gray. SPRING coordinates, cluster labels, pseudotimes and a similar visualization were first reported in Ref. (23). (B)-(D) Observables of neutrophil trajectory as a function of pseudotime calculated for each of 121 bins of pseudotemporally adjacent cells. All bins had 1000 cells except for the last one which had 1310 cells, and had a 50% overlap with neighboring bins. (B) Average gene expression in pseudotemporal bins for highly expressed (max(⟨expr ⟩) *>* 1) and highly varying (coeff. of var.(⟨expr ⟩) *>* 0.5) genes. (C) Largest covariance eigenvalue (black) compared with a statistical null (gray, details in Secn. S3), shifted to have 0 min. Error bars of null are one SD. (D) Distribution of normalized gene expression for each cell projected onto the convariance eigenvector at *τ*_*m*_. Dashed lines indicate bifurcation pseudotimes.

There were several features of this trajectory that make it ideal for applying our analysis framework: it is robust to the systematic, temporal controls of sequencing-time and cellular bar-codes, and included a large number of cells, enabling statistically reliable covariance and correlation measurements. Finally, this trajectory is part of hematopoiesis, a well-characterized developmental process, allowing us to compare our findings against past work.

To determine if the transitions from HSC to neutrophil were due to bifurcations in transcriptomic space, we split the neutrophil trajectory into overlapping bins of 1000 cells (last bin had 1310; details in Secn. S5 and Fig. S5) and applied our covariance analysis to the full, row (cell)-normalized transcriptomic matrix at each bin **G**(*τ*). We found that the largest eigenvalue of the covariance matrix (*ω*_1_(*τ*), dark line in Fig. 4C) exhibited very little variation for *τ < τ*_*d*_ = 85, but began to increase at *τ*_*d*_, and exhibited a significant spike at *τ*_*m*_ = 109. To determine if these *ω*_1_ changes were statistically significant, we computed the corresponding statistical null (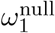, lighter line in Fig. 4(C), details in Secn. S3) and found that the large peak at *τ*_*m*_ was easily distinguishable from the null. We first focus our attention at the dynamics at *τ*_*m*_, following which we will address those observed at *τ*_*d*_. As this pattern of a statistically significant spike following near-constant *ω*_1_ echoed the observed behavior of a saddle-node bifurcation in our toy model (Fig. 3D), we speculated that at *τ*_*m*_ there was a one-to-one transcriptomic state transition. To further visualize the multimodality due to this one-to-one state transition, we computed the projection of normalized gene expression onto the first covariance eigenvector at the bifurcation 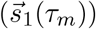. We found (Fig. 4D) that similarly to the saddle-node bifurcation (Fig. 3E) the distribution of this projection was widest at the transition point, *τ*_*m*_, providing further evidence that the cells switched from one transcriptomic state to another at *τ*_*m*_, and that at *τ*_*m*_ the transcriptomic state was multimodal. To verify that this temporal trend was not limited to the diffusion-based pseudotime algorithm used in Ref. (14), we recalculated pseudotime via Slingshot (19), a generic spline-based pseudotime inference tool, and found the same rise and peak of *ω*_1_ (see Secn. S4.2 and Fig. S6A-B).

We focus now on our observations in proximity to *τ*_*d*_. As the increase in *ω*_1_ at *τ*_*d*_ strongly resembled our toy model’s pitchfork bifurcation (Fig. S4B), as well as the proliferation of cell fates seen in high-resolution time-course scRNA-seq experiments (47), we sought to determine if the increase of *ω*_1_ at *τ*_*d*_ was also due to transcriptomic state changes. While 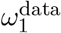 is not obviously well-separated from 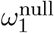 during the increase, the difference between them switched from negative to positive shortly after *τ*_*d*_ (Fig. 4C, inset), indicating a regime crossover. Furthermore, despite gene expression being normalized per cell throughout the dataset, the distribution of each gene’s expression (across cells) begins to significantly shift toward higher values at *τ*_*d*_ (Fig. S5C). These findings suggest that with additional sequencing or prior knowledge, the genomic and geometric details of a one-to-many transcriptomic state transition at *τ*_*d*_ may be distinguishable.

To determine if the transcriptomic state transitions we identified had biological significance, we compared our findings against the tree of cell fates for neutrophil development (Fig. 5A). We found (Fig. 5B) that *τ*_*d*_, the moment *ω*_1_ begins to increase, corresponded well with the moment in pseudotime that cells switched between the endpoints of this tree, from not expressing any terminal-fate marker genes to primarily expressing neutrophil cell-fate markers (23). At a more granular developmental level, our covariance analysis highlights specific transitions between intermediate neutrophil progenitor states (Fig. 5A). These transitions include: (a) one-to-many cell fate changes (i.e., decisions) like the transition between a granulocyte monocyte progenitor (GMP, or myeloblast), and any of its four terminal fates: neutrophil, monocyte, eosonophil, and basophil, as well as (b) one-to-one cell fate changes (i.e., maturation) like the transition between promyelocyte and myelocyte (23, 48, 49). *τ*_*d*_ corresponds well with the point in pseudotime that promeylocyte marker genes are maximal (Fig. 5C and Fig. S6C), suggesting a connection between *τ*_*d*_ and the one-to-many change from GMP to promyelocyte. This can be understood in light of other one-to-many state transitions, like a pitchfork bifurcation (Fig. S4A), where *ω*_1_ increases steadily if different branches are left unclustered (Fig. S4B). Thus, the increase of *ω*_1_ at *τ*_*d*_ suggests that while cells in the neutrophil trajectory were expected to only include the neutrophil lineage branch of GMP, they may in fact include other GMP lineages, such as eosonophils or basophils. Conversely, *τ*_*m*_ correponds well with the point in pseudotime that myelocyte marker genes are maximal, and promyelocyte genes have reduced expression (Fig. 5C and Fig. S6C), suggesting that *τ*_*m*_ denotes the transition point between these two cell fates. This marker-gene analysis matches our geometric interpretation, that an *ω*_1_ spike indicates a one-to-one state transition, as the promyelocyte-to-myelocyte cell fate transition is a maturation step of committed neutrophil progenitors, and not a decision between multiple fates.

**Fig. 5.**
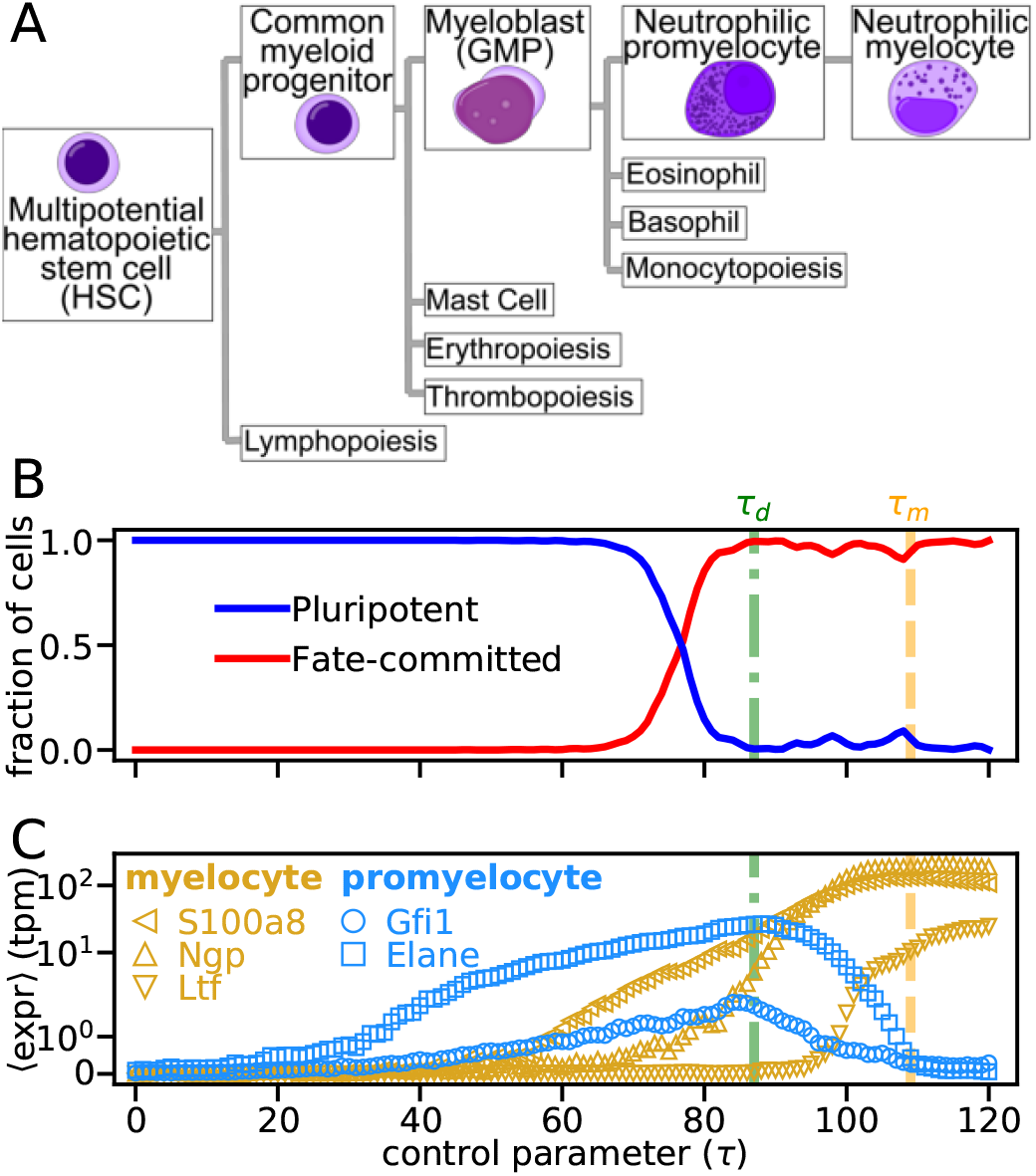
Detected bifurcations correspond to biologically characterized developmental transitions. (A) Schematic of neutrophil development, beginning from hematopoietic stem cells and ending at the neutrophil myelocyte, a committed neutrophil progenitor. Lines indicate naturally occuring progeny, other than the cell type itself. Subsequent neutrophil-committed fates (neutrophil metamyelocyte, band cells, and neutrophils) are not shown. Cell type images by Mikael Häggström, and A. Rad, used with permission. (B) Fraction of cells in each cell type, based on annotated clustering in Ref. (23). (C) Average expression of promyelocyte (blue) and myelocyte (gold) marker genes (23). SEM error bars are smaller than symbols.

Thus, by using Eq. (5) to quantify the geometry of neutrophil development, we were able to recover the known GMP-neutrophil cell fate decision, qualify the trajectory as likely including other lineages, and pinpoint a maturation step in neutrophil development. Broadly, these results qualify the notions portrayed in Waddington’s landscape, as they show that not only do cell fate decisions correspond to (one-to-many) transcriptomic state changes, but that even maturation fate changes can correspond to discontinous transitions between transcriptomic states. Interestingly, the state changes at both *τ*_*m*_ and *τ*_*d*_ appear to occur in similar directions, as revealed by the fact that the distribution of gene expression projected onto 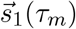 begins to widen at *τ*_*d*_ (Fig. 4D), suggesting that there may be generic transcriptomic modes that more readily enable state changes. Because the promyelocyte-to-myelocyte transition identified at *τ*_*m*_ is distinct from its null, we use this bifurcation to further test the our mathematical framework (Eq. (7)) and present new hypotheses for mechanistic aspects of neutrophil development in mice.

### C. Correlation analysis identifies genetic contributor in myeloid differentiation

As correlation analysis at a bifurcation correctly ranked important gene relationships in our example saddle-node bifurcation (Fig. 3G-H), and has been shown to reveal gene-pair relationships that are critical for state transitions in temporal RNA-seq data (26, 40), we used *R*_*ij*_(*τ*_*m*_), the Pearson’s correlation coefficient between genes *i* and *j* to identify pairwise gene relationships that are important for the promyelocyte-to-myelocyte transition. We verified that the distribution of *R*_*ij*_ widened at *τ*_*m*_, relative to nearby *τ* (Fig. 6A; see Secn. S5 for PCC calculation details) as predicted for one-to-one state transitions (Eq. (7)) and observed in the saddle-node bifurcation example (Fig. 3G)).

We found that the genes in the tails of the *R*_*ij*_(*τ*_*m*_) distribution (details in Secn. S5) formed a well connected correlation network, comprising only 2 connected components (Fig. 6B). By clustering the genes based on their various cellular functions, we determined that the larger component (28/31 genes) showed features of a developmental gene regulatory network. The genes in this cluster included markers that broadly classify neutrophils, as well as markers that more specifically classify neutrophilic promyelocytes, some of which (S100a6, S100a8, S100a9, and Ngp) had previously been used to classify these cells (23), and others (Sirpa, Ccl6) which had not. In particular, the S100a9 gene was found to be highly negatively-correlated with the majority of other genes in the network, suggesting that it is pivotal for the observed bifurcation. This cluster also included genes involved in the mechanisms that drive fate changes, such as signaling and transmembrane processes necessary for cell-cell interactions, as well as housekeeping and metabolic genes responsible for maintaining basic cellular processes. Interestingly, the smaller component (3/31 genes) exclusively comprised mitochondria regulators, indicating that background, highly covariable expression relationships may be difficult to filter out exclusively from correlation matrix analysis (Fig. 6B). Thus, bifurcation detection enables inferring gene regulatory networks for a specific moment in development from a physically principled, dynamical systems approach.

**Fig. 6.**
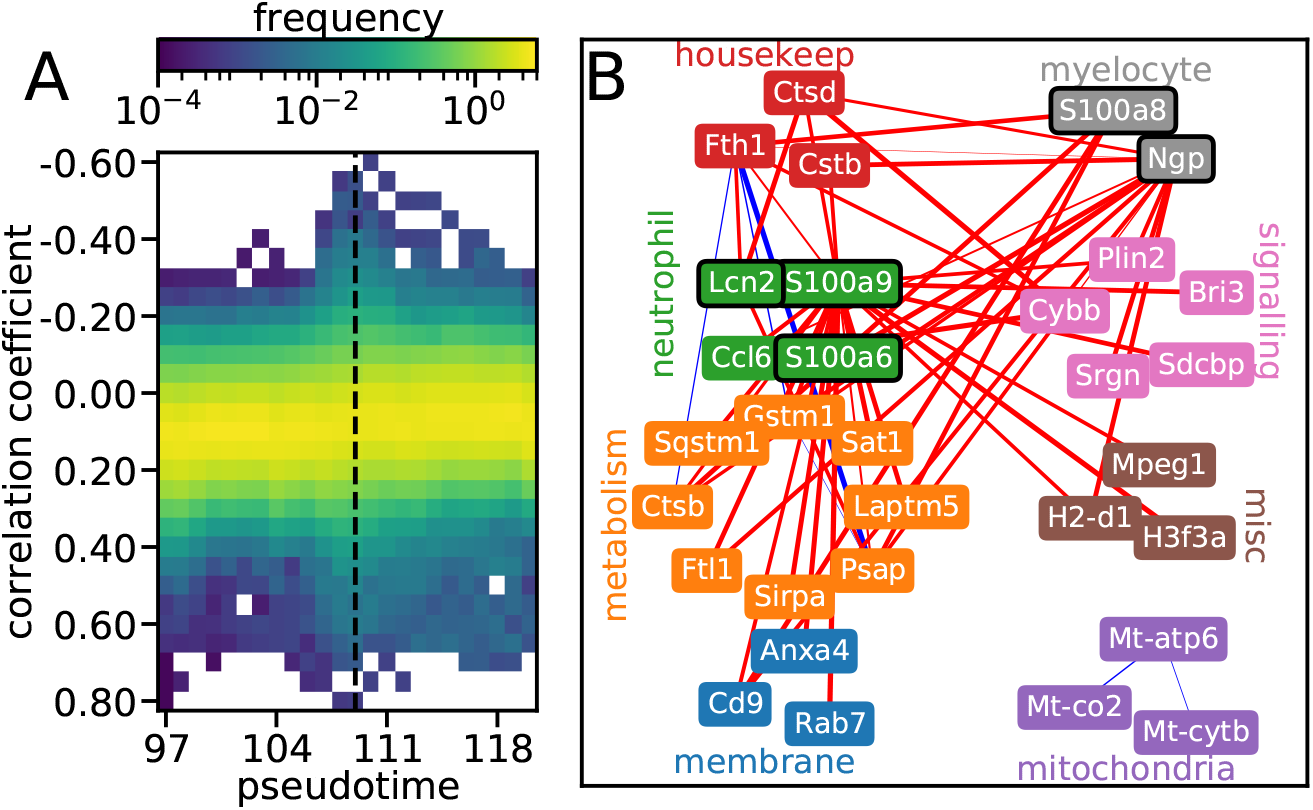
Correlation analysis at bifurcation reveals critical genes for developmental transition. (A) Distribution of correlation coefficients between highly coincidentally expressed gene pairs (details in Secn. S5) as function of pseudotime. Columns integrate to 1. Dashed line indicates bifurcation bin. (B) Correlation graph for gene pairs that have *R*_*ij*_ (*τ*_*m*_) *<* −0.35 or *R*_*ij*_ (*τ*_*m*_) *>* 0.65, grouped by cellular function. Edge color indicates positive correlation (blue) or negative (red) and thickness is proportional to correlation magnitude (separate scale for positive and negative correlation.)

We also found that the gene expression patterns within the clusters reflected their importance in the promyelocyte-to-myelocyte transition (Fig. S7). Neutrophil and myelocyte cell fate markers, as well as genes implicated in signaling, appeared to switch between low and high expression at *τ*_*m*_, indicating that they have expression patterns tied to specific cell fates. Conversely, metabolic, membrane, and housekeeping genes nearly exclusively exhibit local peaks in expression at *τ*_*m*_, indicating that they are transiently important for driving, or guiding, the cell-fate transition but do not pertain to a specific fate. The mitochondrial cluster, as well as some miscellaneous and membrane genes (H2d1, Anxa4) exhibited gradual increases or decreases in expression, indicating that they may have developmental patterns that are generically tied to cell fate specification, or distance from pluripotency, but not specifically to the promyelocyte-myelocyte transition.

Lastly, since the principal covariance eigenvector 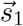 is equivalent to the soft mode of the Jacobian at a bifurcation (Eq. (6) and Fig. 3F), we examined how the 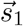 loadings of the network genes varied around *τ*_*m*_ to determine if they implied a deeper mechanistic interpretation (Fig. S8). We found that for nearly all genes in the developmental cluster, loadings were locally maximal at *τ*_*m*_, suggesting that their relationships with other genes are most important at *τ*_*m*_, even if their absolute expression is tied to a specific fate (e.g., neutrophil markers). Interestingly, some marker genes (Ngp, S100a8, S100a9) appeared to transition between high and low loading values at *τ*_*m*_, indicating that they may maintain the specific cell fates of promyelocyte or myelocyte through their interactions with other genes. The loadings also appeared to reiterate the lack of importance of genes in the mitochondrial cluster for the promyelocyte-to-myelocyte fate change, as their 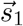 loadings were independent of pseudotime.

In summary, by applying our mathematical framework, that genetic relationships closely tied to a fate change would be highly correlated at a bifurcation (Eq. (7)), and that a bifurcation’s principal eigenvector is mechanistically informative (Eq. (6)) we recovered genes that were known to be important in neutrophil development for mice, and could place them in context of a larger, newly hypothesized gene regulatory network in data-driven manner. We verified the importance of many of these genes by examining their expression in a pseudotime window around the promyelocyte-to-myelocyte fate change, and are thus able to predict the extent to which individual genes are either transiently important for a cell fate transition, or characteristic of a specific cell fate. The eigenvector loadings were particularly useful in determining whether gene pairs are highly correlated at a bifurcation because of their importance to other genes, maintaining specific cell fates, or pluripotency. Thus, we claim that the mathematical framework presented here not only helps elucidate the dynamics of the transcriptome at the scale of a cell’s fate, but even provides clues at the scale of molecular interactions for the underlying genetic relationships that enable cell-scale changes.

## 4. Discussion

It is now both appealing and common place to loosely graft dynamical systems language to the dynamics of cellular differentiation. In particular, it is appealing to allude to bifurcations of an underlying landscapes as driving qualitative transitions in accessible states. But what are the necessary statistical signatures that must be present in transcriptomic trajectory data such that a bifurcating dynamical systems point of view is an accurate description? And, if such signatures are present, what further theory-based quantitative insights can be had that go beyond word-based analogies?

One of the foundational principles of the study of bifurcations is the concept of linear stability, and its loss. The theory outlines that a generic way for a stable solution of the steady-state equations of motion to lose stability is through the change of sign of an eigenvalue/s of the Jacobian. In this context, the system is “free” in the direction of the corresponding eigenvector/s. Naturally, the system isn’t unbounded, and non-linearities emerge to stabilize a new stable state of the system. In this study, we show that covariances of the state-variables are sufficient to assess whether linear stability is being lost, through leveraging the continuous-time Lyapunov equation. Our central result, based on the analysis of a restricted portion of the transcriptomic trajectories present during the process of hematopoiesis, is that the requisite statistical signatures of a bifurcation through a loss of linear stability are detectable and present during development. Once detected, additional features, such as the unstable eigenvector and the correlations between the state-variables can be identified. From a data-analysis point of view we demonstrate a linear statistical treatment of pseudotime-based transcriptomic trajectories, which is both computationally trivial and statistically robust. As the ability to construct increasingly accurate trajectories improves, our methodology should only become more robust and precise at characterizing critical moments in development. From a more theoretical point of view, our work takes a step toward inferring mathematical models for key developmental decisions at a whole genome scale to mirror the several beautiful dynamical systems biology inferred from imaging studies (43)[NEED CITATIONS].

Our finding that a transcriptomic trajectory can have distinct geometric signatures, including durations during which the principal covariance eigenvalue is constant or spikes, has considerable consequences for understanding scRNA-seq datasets that capture developmental transitions. That we saw any consistent behavior in the principal covariance eigenvalue lends significant support to our initial assumption that cell fate modifiers operate at a much slower rate than transcriptomic modifiers, because if these occured on similar timescales, no statistical signature would be evident, let alone those that align well with our current developmental understanding of the system. Additionally, while previous statistical analyses of scRNA-seq data found that developmental trajectories appear as monotonic proliferations of cell fates (47), our focus on a single developmental trajectory enabled distinguishing multiple developmental epochs, including durations of development during which cell fates did not undergo qualitative changes, proliferated (the GMP-to-promyelocyte transition), and changed state (the promyelocyte-to-myelocyte transition). Our evidence of bifurcations also starkly contrasts with scRNA-seq visualizations that show gene expression varying smoothly along a developmental path, and underscores the importance of understanding noise and non-linear dependencies when using transcriptomic profiles to classify a cell’s fate (17).

While our results have been derived using bifurcation theory, such that the fixed-point stability for developmental dynamical systems changes as a function of developmental time, they do not preclude the possibility that cell-fate transitions may occur due to stochasticity (15, 38). To explore if stochasticity may have contributed to neutrophil development, we simulated a noise-induced transition using the same model gene regulatory network (see Secn. S6 for details). We found that the principal covariance eigenvalue exhibited step-like dynamics as the stochasticity was varied (Fig. S9), unlike either the one-to-one (Fig. 4C) or one-to-many (Fig. S4B) bifurcation examples previously explored. As this step-like signature for the principal covariance eigenvalue did not resemble the steady increase and spike in the neutrophil trajectory (Fig. 4C), while bifurcations did, our results suggest that neutrophil differentiation likely occurs due to a loss of stability between fixed points, rather than noise-induced transitions.

Our analysis of the data in Ref. (23) also yielded intriguing implications regarding the specific geometry of neutrophil development in mice. In particular, some of the known cell fate changes in neutrophil development were not distinguishable in the covariance eigenvalue trajectory (e.g. from Common myeloid progenitors [CMP] to GMP), which indicates that these changes are less discontinuous than the GMP-to-promyelocyte or promyelocyte-to-myelocyte transitions. This could mean, for example, that even when CMPs differentiate to GMPs, the transition lacks committment, and is dependent on a sustained developmental signal, whereas once cells transition from GMP to promyelocyte, they are committed to becoming neutrophils regardless of an external signal. Furthermore, in projecting gene expression onto the principal covariance eigenvector at the promyelocyte-to-myelocyte fate transition (Fig. 4E), it became apparent that direction along which that fate change happened was well aligned with the direction of the GMP-to-promyelocyte transition. This result may be a sign of distinct, soft directions in transcriptomic space along which cell fates are most likely to change.

Aside from these geometric implications, pinpointing the bifurcation in pseudotime may also enable efficent identification of the genes and molecular mechanisms that drive a cell fate transition. As shown, important pairwise gene relationships are explicitly highlighted in the correlation matrix at a bifurcation. While these correlations lack directionality, and therefore cannot be used to infer all aspects of mechanism, they may aid in building regulatory network models when combined with prior protein-interaction data or new experimental perturbations (50). Additionally, since the principal covariance eigenvector is equivalent to the Jacobian eigenvector at the bifurcation, and the Jacobian directly reflects gene dynamics, the eigenvector may reveal critical dynamic information or constrain an inferred global Jacobian (51).

While we focus here on scRNA-seq data, our approach is broadly applicable, and could, in principle, aid in determining other aspects of developmental bifurcations, such as the genomic structural modifications necessary for fate transitions from single-cell ATAC-seq data (52, 53). A particularly fascinating direction to pursue in the future is a comparative study of developmentally and/or evolutionary related trajectories and the nature of modifications of the bifurcations. Additionally, it may be possible to incorporate our covariance analysis into other indications of pseudotime rank, such as cellular barcodes and low-dimensional distance, to constrain developmental trajectories along bifurcative paths. Our analysis was only possible because scRNA-seq experiments can now measure the expression of tens of thousands of genes in hundreds of thousands of cells, enabling accurate covariance measurements. That we found bifurcative events in this data implies that there are low dimensional, non-linear dynamical systems at play, and that sufficient biological sampling will reveal the knobs to controllably tilt developmental landscapes.

## Materials and Methods

Instructions and Python code for reproducing all figures in this manuscript are available at github.com/simfreed/sc_bifurc_figs.

## 5. Derivation of results in Secn. 2

### A. Continuous time Lyapunov equation for transcriptomic matrices

Let **G** be the steady state transcriptomic matrix at a single developmental time with *n*_*c*_ rows (cells) and *n*_*g*_ columns (genes) (Fig. 2A) and **F** be a set of differential equations describing the molecular interactions that generate **G**, such that

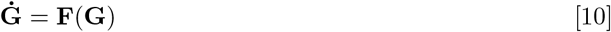

where **Ġ** is the derivative of **G** with respect to time. Since all cells (columns) in **G** are at steady state at the same developmental time *τ*, we assume (for the purpose of contradiction) that they are all statistical replicates of the same transcriptomic state, 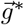, and the full matrix, **G**, is therefore in the vicinity of the hyperbolic fixed point

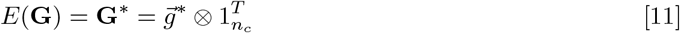

where 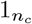 is a vector of *n*_*c*_ ones and *E* denotes the expectation operator. The dynamics of **G** can be by linearized via the distance to the fixed point **X** = **G** − **G**\*, such that

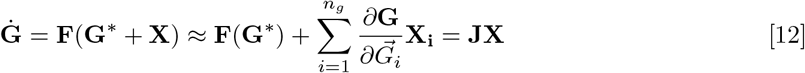

where 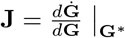 is the Jacobian of **G** and we have used the fact that at steady state, **F**(**G**\*) = 0.

If **F** is stochastic and Markovian, then the dynamics of **X** can be described as a discretized Ornstein-Uhlenbeck (OU) process,

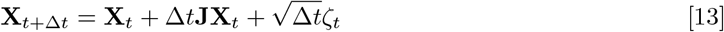

where Δ*t* is the molecular interaction timescale and *ζ*_*t*;*i,j*_ is sampled from *N* (0, *σ*_*i*_) where *σ*_*i*_ is the variance of gene *i*. The gene-gene covariance matrix can then be defined as

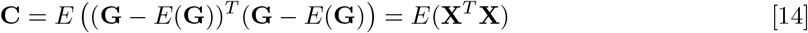

where the superscript *T* denotes transpose and we have approximated *E*(**G**) = **G**\*. The stationary condition for an OU process, (i.e., that *∂***C***/∂t* = 0) then yields

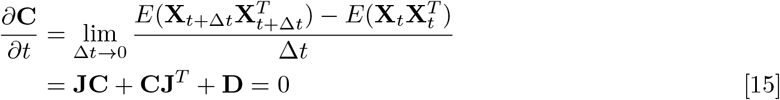

where 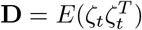 and we have used the fact that *E*(*ζ*_*t*_),*E*(**X**_*t*_),*E*(*ζ*_*t*_**X**_*t*_), and *E*(**X**^*T*^ *ζ*^*T*^) are all 0 (31).

### B. Covariance at bifurcation

If **J** is diagonalizable, such that,

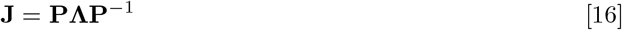

where Λ is a diagonal matrix of eigenvalues 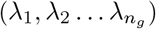 and **P**^*T*^ is the square matrix of eigenvectors 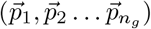, then Eq. (15), often referred to as the continuous-time Lyapunov (CL) equation, can be used to qualitatively assess **G**. Left-multiplying Eq. (15) by **P**^−1^ and right-multiplying by (**P**^†^)^−1^, yields

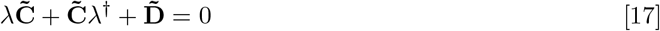

where the † superscript indicates conjugate transpose, 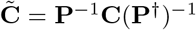 and 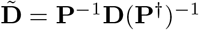. Since **Λ** is diagonal, Eq. (17) can be rewritten elementwise,

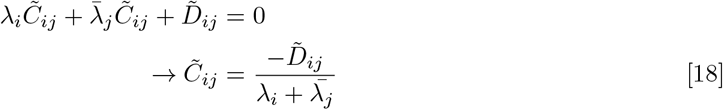

which can be substituted to yield an expression for elements of the covariance

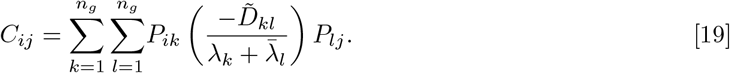

since 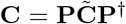 At a bifurcation, max (**Λ**) = *λ*_*d*_ → 0, so the *k* = *l* = *d* term in Eq. (19) becomes dominant, and

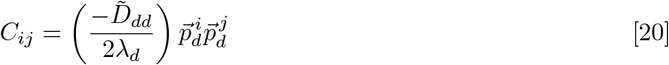

where 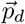 is the *d*^*th*^ column of **P**.

### C. Bifurcation eigenvector equivalence

Since **C** is real and symmetric, the eigenvalue decomposition can be written as a single sum,

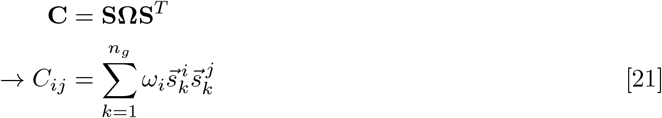

where 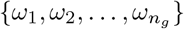 are its eigenvalues, and 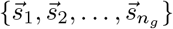 are its eigenvectors, which are normalized to 1. For large *n*_*g*_, **Ω** and **S** are signficantly easier to compute than **C** as they can be obtained from the singular value decomposition of **X** = **G** − *E*(**G**). For Eq. (21) to be equivalent to Eq. (20) at a bifurcation, at least one eigenvalue *ω*_*i*_→ ∞, which we may, without loss of generality, refer to as *ω*_1_. If *ω*_1_ ≫*ω*_*i*_ for *i* ∈ [2 … *n*_*g*_] then the *k* = 1 dominates the sum in Eq. (21), and by equating with Eq. (20) we obtain

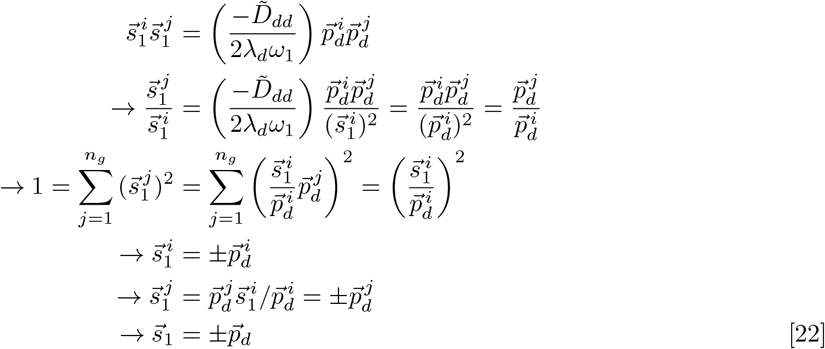

where we have used the fact that 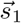 and 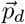 both normalize to 1.

## 6. Simulation methodology

To explore our analysis framework on a more biologically relevant gene network (Fig. 3A) we utilized a Focker-Plank simulation method. For each of the *N*_*c*_ = 100 cells (*N*_*c*_ chosen by examining how many cells were neccesary to accurately detect bifurcations in the neutrophil data Fig. S5A-B) the expression of gene *i*, (*g*_*i*_(*t*; *m*_1_, *m*_2_, *k*_*D*_) is initalized uniformly randomly in the interval (0,4]. The expression at subsequent timesteps (*g*_*i*_(*t* + Δ*t*)) is sampled from a Gaussian distribution *N* (*μ, σ*) where

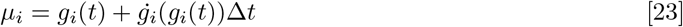

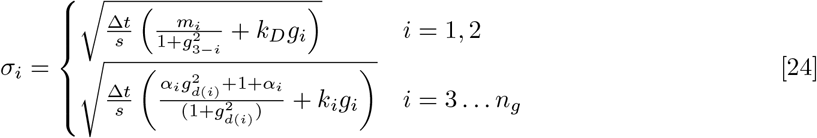

and bounded to be non-negative. The simulation ran for *N*_*t*_ = 1*e*7 timesteps, with Δ*t* = 0.01 as that was sufficient for equilibration Fig. 3, at a noise scale of 1*/s* = 0.05, and the last timestep of each simulation is the steady state expression **G**. In the saddle-node example (Fig. 3) the remaining parameters were *k*_*D*_ = 1, *m*_2_ = 3, *m*_1_ ∈ [2, 4] while in the pitchfork example (Fig. S4) *m*_1,2_ = 1, *k*_*D*_ ∈ [0.24, 5].

## 7. Analysis pipeline

Eq. (5)-7 imply an analysis pipeline characterizing bifurcations in high-dimensional temporal data, which we use in this paper:

1. Obtain highly sampled temporal data **Caveat:** for data types such as scRNA-seq, where frequent sampling is difficult, and samples may include realizations from many different times, time may be inferrable, using, for example pseudotime inference (see Secn. S4.1)
2. Bin the data along the temporal axis (see Secn. S5).
3. Compute the largest eigenvalue of the covariance matrix (*ω*_1_) in each bin (e.g., using an off-the-shelf PCA function).
4. Evaluate if a bifurcation occurs by comparing *ω*_1_ with a suitable null (see Secn. S3).
  a. Spike indicates a one-to-one bifurcation.
  b. Steady increase indicates a one-to-many bifurcation.
5. If a bifurcation is detected (e.g., at *τ*_*c*_) compute and examine the correlation matrix **R**(*τ*_*c*_), as entries of **R**(*τ*_*c*_) that are large in magnitude are critical to the bifurcation at *τ*_*c*_.

## ACKNOWLEDGMENTS

We thank Richard Carthew, Yogesh Goyal, and Karna Gowda for reviewing the manuscript and providing suggestions. This work was supported by the NSF-Simons Center for Quantitative Biology at Northwestern University and Simons Foundation (597491, MM). SG acknowledges University of Toronto’s Medicine by Design initiative, which receives funding from the Canada First Research Excellence Fund, and NSERC Discovery Grant MM is a Simons Foundation Investigator.

## 8. Supporting Information

### S1. Methodological relationship to Dynamical Network Biomarkers

Chen et al. (25), developed the concept of a dynamical network biomarker (DNB), a group of genes that drive a critical transition and are detectable from high dimensional gene expression datasets. In particular, they define an indicator function

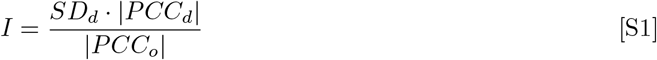

where *SD*_*d*_ is the average standard deviation of genes in the DNB, *PCC*_*d*_ is the average correlation coefficient between genes in the DNB, and *PCC*_*o*_ is the average correlation coefficient between genes in the DNB and genes outside the DNB (25). At a critical state transition, or bifurcation, *I* is predicted to diverge, because *SD*_*d*_ and |*PCC*_*d*_ |become large, while |*PCC*_*o*_| becomes small. Mathematically, the genes in the DNB correspond to those that have non-zero weight in the direction of the transition, i.e., 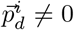, where 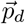 is the principal eigenvector of the Jacobian, while genes outside of the DNB have 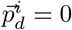. This prediction is qualitatively similar, but not the same as Eq. (5). In particular, while both *SD*_*d*_ and *ω*_1_ increase at a bifurcation, they are not equivalent, as *SD*_*d*_ measures the variance of each individual gene, while *ω*_1_ measures the variance across all genes, and therefore accounts for corrections to the total variance due to covariances between genes in the network. Therefore, for bifurcation detection, we focus solely on *ω*_1_, instead of incorporating correlations into the indicator as in Eq. (S1).

As for determining which gene relationships are critical for the bifurcation, we take a similar approach to Refs. (25, 26), in focusing on the correlations that approach ±1 at the bifurcation. This is justified via Eq. (7), which yields that *R*_*i,j*_ → ±1 if 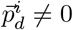 and 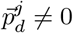. Interestingly, while we derived Eq. (7) via the eigendecomposition of the covariance matrix, Refs. (25, 26) derived the same result form the covariance matrix itself, providing additional support to this method.

### S2. Bifurcations possibilities from two mutually inhibiting genes

At steady state, Eq. (8) satisfies the quintic polynomial

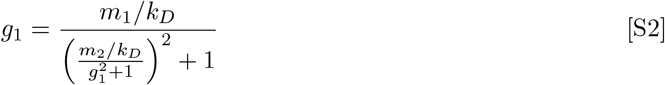

which, depending on the parameter values, can have one real solution that is an attractor (e.g., if *m*_1,2_ = 1 and *k*_*D*_ = 1) or three real solutions, two attractors (nodes) and one repellor (saddle) (e.g., *m*_1,2_ = 1, *k*_*D*_ = 1*/*3). By examining the null clines,

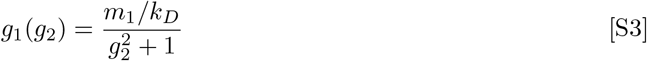

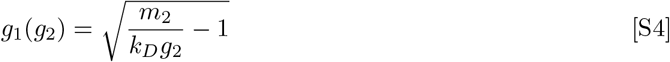

it can be deduced that varying *m*_1_, while fixing *τ* and *m*_2_ can yield a saddle-node bifurcation, as Eq. (S3) moves vertically while Eq. (S4) does not, allowing for either node to merge with the saddle (Fig. S1A). Conversely, varying *k*_*D*_, while fixing *m*_1,2_ and *m*_2_, can yield a pitchfork bifurcation, as both null clines move, such that above the bifurcation value, all three real solutions remain (Fig. S1B). Solving Eq. (S2) computationally via the Python function numpy.roots and plotting the real solutions (Fig. S1C-D) yields the bifurcations used in Fig. 3 and Fig. S4 (54).

### S3. Resampling principal eigenvalue

Given the transcriptomic matrix 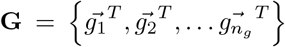, where 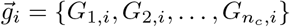 and *G*_*i,j*_ is the expression of the *j*^*th*^ gene in the *i*^*th*^ cell, we generate a null sample **G**^null^ by drawing each of its entries 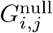 randomly, with replacement, from 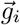. In Fig. S5,3,4, we compute the principal covariance eigenvalue 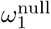 for each of *n*_*s*_ = 20 samples, and compare this null distribution against the principal covariance eigenvalue of the data *ω*_1_. This resampling technique has little impact on *ω*_1_ for unimodal distributions as the scale of *ω*_1_ is still determined by the system’s noise (Fig. S3 left and right), but significantly decreases *ω*_1_ for multimodal distributions (Fig. S3 center) since the structure of the multimodality is scrambled; thus we found it was an effective method for determining if a spike in *ω*_1_ is due to multimodality or increased noise.

**Fig. S1.**
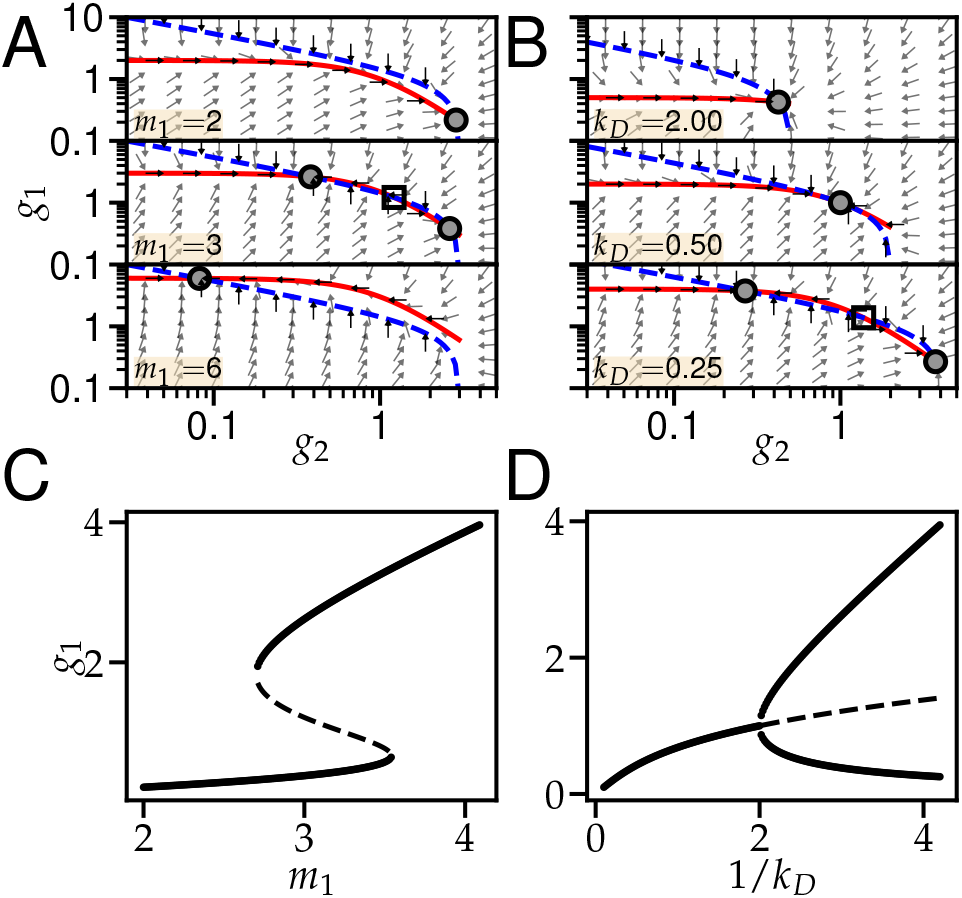
Analysis of Eq. (8). (A-B) Phase planes for different parameter sets yield a saddle-node bifurcation (A) or pitchfork bifurcation (B). Solid red line is given by Eq. (S3) while dashed blue line is given by Eq. (S4). Open squares are saddles while closed circles are nodes. Arrow angles are given by *tan*^−1^(*ġ*_1_*/ġ*_2_) and are uniform length. (C-D) Solutions to Eq. (S2) while varying *m*_1_ (C) or *k*_*D*_ (D). Solid lines are nodes and dashed lines are saddles. In (A,C) *k*_*D*_ = 1, *m*_2_ = 3 and in (B,D) *m*_1,2_ = 1.

### S4. Pseudotime inference

#### S4.1. Algorithm for generating the pseudotime labels in Weinreb et al

SPRING (x-y) positions, cell type annotations, and pseudotime ranks for the data presented in Fig. 4A-B were downloaded from https://github.com/AllonKleinLab/paper-data/tree/master/Lineage_tracing_on_transcriptional_landscapes_links_state_to_fate_during_differentiation. The algorithms to generate these values are described in detail in Ref.(23)(Supplementary Materials) and recapitulated here for completeness. Given the full *in-vitro* hematopoiesis transcriptomic matrix (all cells and all genes), the SPRING positions in Fig. 4A plot were generated using the following procedure.

1. A filtered transcriptomic matrix was generated which did not include genes that
  a. had low variability as determined via the filter_genes function with parameters (85,3,3) from https://github.com/AllonKleinLab/SPRING_dev/blob/master/data_prep/spring_helper.py (14).
  b. correlated highly (R>0.1) across all cells with any of the following cell cycle genes: Ube2c, Hmgb2, Hmgn2, Tuba1b,Ccnb1, Tubb5, Top2a, and Tubb4b.
2. The top 50 principal components (PC) of the filtered transcriptomic matrix were computed.
3. 40,000 of the cells were selected randomly, and a k-nearest-neighbors (KNN) graph between those cells was constructed using the top 50 PC of the filtered transcriptomic matrix and k=4.

**Fig. S2.**
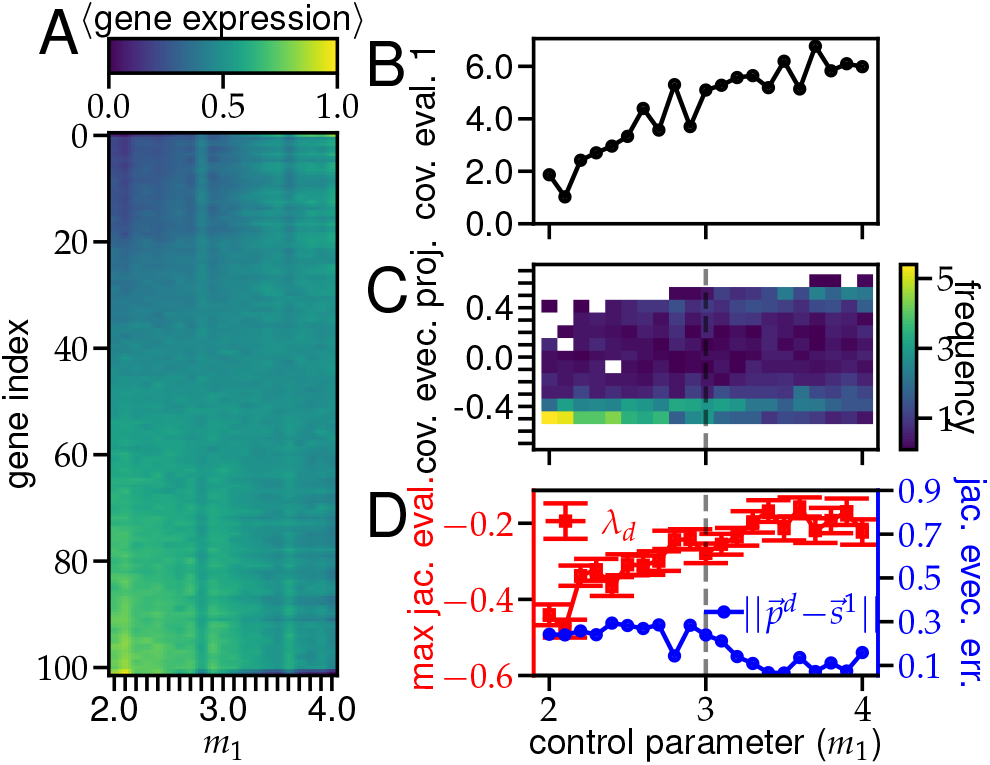
Unequilibrated saddle-node bifurcation analysis. (A)-(D) Corresponding plots for Fig. 3 (C)-(F), respectively, when the simulation is runs for a *N*_*t*_ = 500 iterations rather than *N*_*t*_ = 50000.
4. X-Y positions of these 40,000 cells were generated using the ForceAtlas2 algorithm with 500 steps (55).
5. Positions for each of the remaining 90,887 cells were computed as the average position of their 40 nearest neighbors (in the 50-PC space) among the initial 40,000 cells.

Cells were annotated with their cell types (cluster annotation in Fig. 4A) based on their position in the SPRING plot and their expression (terminal cell fates) or lack of expression (pluripotent) of pre-selected marker genes. Specifically the marker genes used to determine if cells were neutrophils were S100a9, Itgb2l, Elane, Fcnb, Mpo, Prtn3, S100a6, S100a8, Lcn2, and Lrg1.

Neutrophil pseudotime rank was then determined by smoothly interpolating between cells in the pluripotent and neutrophil clusters. The interpolation method used throughout this procedure is an iterative, diffusive process defined as

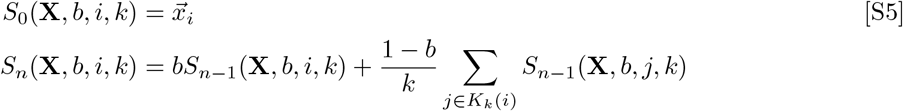

where 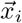 is a vector quantity defined for cell 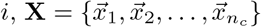 is the matrix of this quantity for all cells, *K*_*k*_(*i*) are the cell indices of the *k* nearest neighbors of cell *i, n >* 0 is the number of iterations, and *b* is the neighbor weight (low *b* and high *n* both yield high diffusion) (11). The pseudotime ranking procedure is:

1. Cells are identified to be part of the neutrophil trajectory
  a. Let 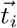 be an indicator vector for the cell type of *i*; i.e. *t*_*ij*_ = 1 if cell *i* is type *j* and 0 otherwise. Let 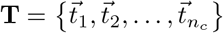 be the corresponding matrix for all cells.
  b. Let **K**_100_ be the k-nearest-neighbor graph between cells for *k* = 100 using the top 50 PC.
  c. Let 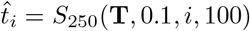 be the smooth cell type indicator.

**Fig. S3.**
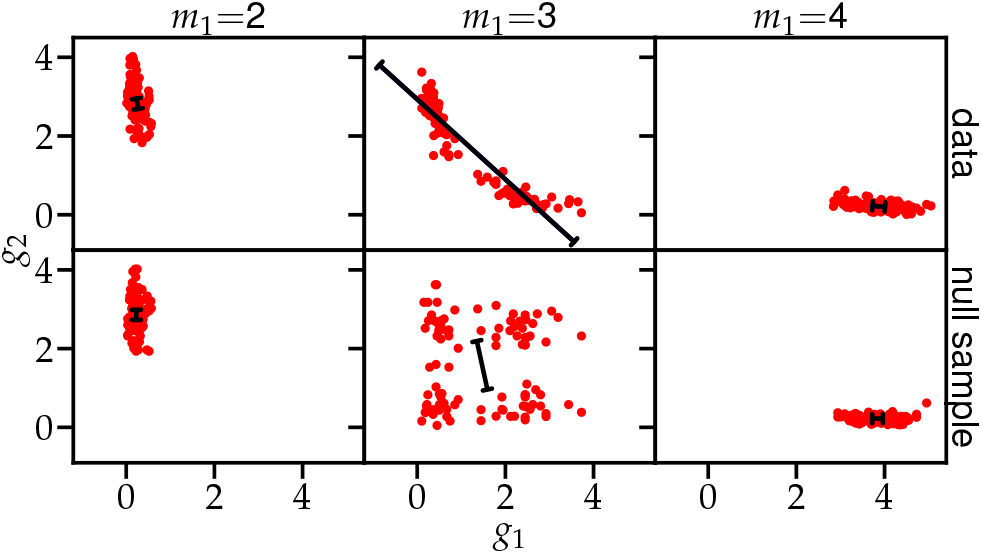
Effect of resampling on principal eigenvector. Red dots are cells for the data (top) and a single marginal resampling (bottom) before the bifurcation (left), at the bifurcation (center) and after the bifurcation (right). Direction of the black lines corresponds to the principal eigenvector and length corresponds to the principal eigenvalue.
  d. Let 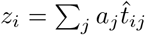 be the weighted average cell type 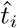 where the weights for each cell type (*j*) are

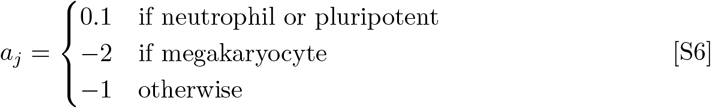
  e. Let 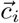 be a neutrophil trajectory indicator such that 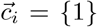 if *z*_*i*_ > *Q*_0.6_(*z*) and {0} otherwise, where *Q*_0.6_(*z*) is the 60^th^ quantile of *z*. Let 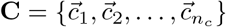.
  f. Let *ĉ*_*i*_ = *S*_50_(**C**, 0.1, *i*, 100) be the smoothed neutrophil trajectory indicator.
  g. Cells were considered part of the neutrophil trajectory if *ĉ*_*i*_ *> Q*_0.6_(*ĉ*) where *Q*_0.6_(*ĉ*) is the 60^th^ percentile of *ĉ*.
2. The 61, 310 cells identified as part of the neutrophil trajectory are sorted
  a. Let 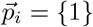 if a cell in the trajectory is pluripotent and 0 otherwise; i.e., it is an indicator for pluripotency. 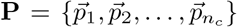 is the corresponding matrix for all cells in the trajectory.
  b. Let 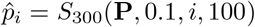 be the smoothed pluripotency indicator.
  c. The pseudotime of cell *i* is the rank (largest to smallest) of 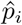 among all 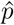.

#### S4.2. Pseudotime inferrence in the absence of metadata

To test if the neutrophil bifurcation characterization was dependent on the choice of pseudotime algorithm, we used the Slingshot algorithm (19) to compute the pseudotime of each cell for its trajectory from the undifferentiated cluster to each of the terminal fate clusters. The input to Slingshot were the cells’ cluster labels and their SPRING coordinates, and the output was a probability, or weight, that a cell belonged to each undifferentiatedto-terminal-fate trajectory, as well as its pseudotime along that trajectory. In Fig. S6A, we show the pseudotime of all cells that had *weight >* 0 for belonging to the trajectory that led from undifferentiated cells toward neutrophils. Unlike the pseudotime method described in Secn. S4.1, the origin of the trajectory does not coincide with the earliest sequenced cells, as time of sequencing and clonal barcode data could not be input to Slingshot. Nevertheless, we obtain a clear bifurcation signature in the principal covariance eigenvalue (Fig. S6B) at the point where promyelocyte gene expression decreases to 0 and myelocyte marker gene expression become maximal (Fig. S6C). This result supports our belief that the bifurcation characterization does not depend on the specific pseudotime calculation.

### S5. Computing correlation coefficients

A significant challenge in accurately measuring *R*_*ij*_(*τ*), the Pearson’s correlation coefficient between genes *i* and *j* at pseudotime *τ* (Fig. 6) is that the low readdepth of scRNA-seq experiments results in many genes having 0 or very few reads in many cells, which leads to systematically high correlations (e.g., if the read counts for both gene *i* and *j* are almost all 0, but there is one cell in which both are highly expressed). To mitigate this issue, we: (a) used large bins of 1000 or more cells for all experimental calculations (even though smaller bins are sufficient to detect bifurcations, as shown in Fig. S5A-B), (b) only use cells that have non-zero read count of both genes *i* and *j* (effectively viewing 0 reads as a lack of information, rather than a measurement) and (c) only calculate correlations between gene pairs that had 400 or more non-zero read count cells to ensure that correlations are not spuriously increased by low read counts. While this filter yields a significant reduction in correlations measured (an average of 5.5 * 10^4^ per pseudotime window, out of a possible 3.2* 10^8^), it is independent of pseudotime, and therefore, with respect to the widening of the *R*_*ij*_(*τ*) distribution at *τ*_*m*_ (Fig. 6A), can be thought of as a random statistical sample of correlation coefficients for which there is high confidence. Genes used to form the network in Fig. 6B had *R*_*ij*_(*τ*_*m*_) *>* 0.65, or *R*_*ij*_(*τ*_*m*_) *<* −0.35 as determined via inspection of Fig. 6A.

### S6. Noise induced transitions

To determine if a non-bifurcating noise-induced transition model (15, 38) could yield a similar covariance eigenvalue signature to our observations in the neutrophil trajectory (Fig. 4C), we ran the 100 gene network model (Fig. 3A) in a regime of the dynamical system that had two fixed-points (*m*_1,2_ = 1, *k*_*D*_ = 1*/*3) at varying noise scales *s* (see Fig. S1 and Eq. (24) for details). To ensure a transition, we initialized all cells to populate the fixed point with higher *g*_1_. We found that for low noise values (1*/s* ≤ 0.01) the cells stayed near their initial fixed point, yielding a unimodal distribution for *g*_1_ (Fig. S9A) and low principal covariance eigenvalue (Fig. S9B) while for high noise values (1*/s* ≥ 0.02) the cells visited both fixed points, yielding a bimodal distribution for *g*_1_, and a high principal covariance eigenvalue. As the covariance eigenvalue signature of the neutrophil trajectory did not include these switch-like dynamics, we suggest that in the neutrophil trajectory, the state transitions result from the underlying dynamical systems bifurcating rather than the scale of biological noise overcoming a fixed potential barrier.

**Fig. S4.**
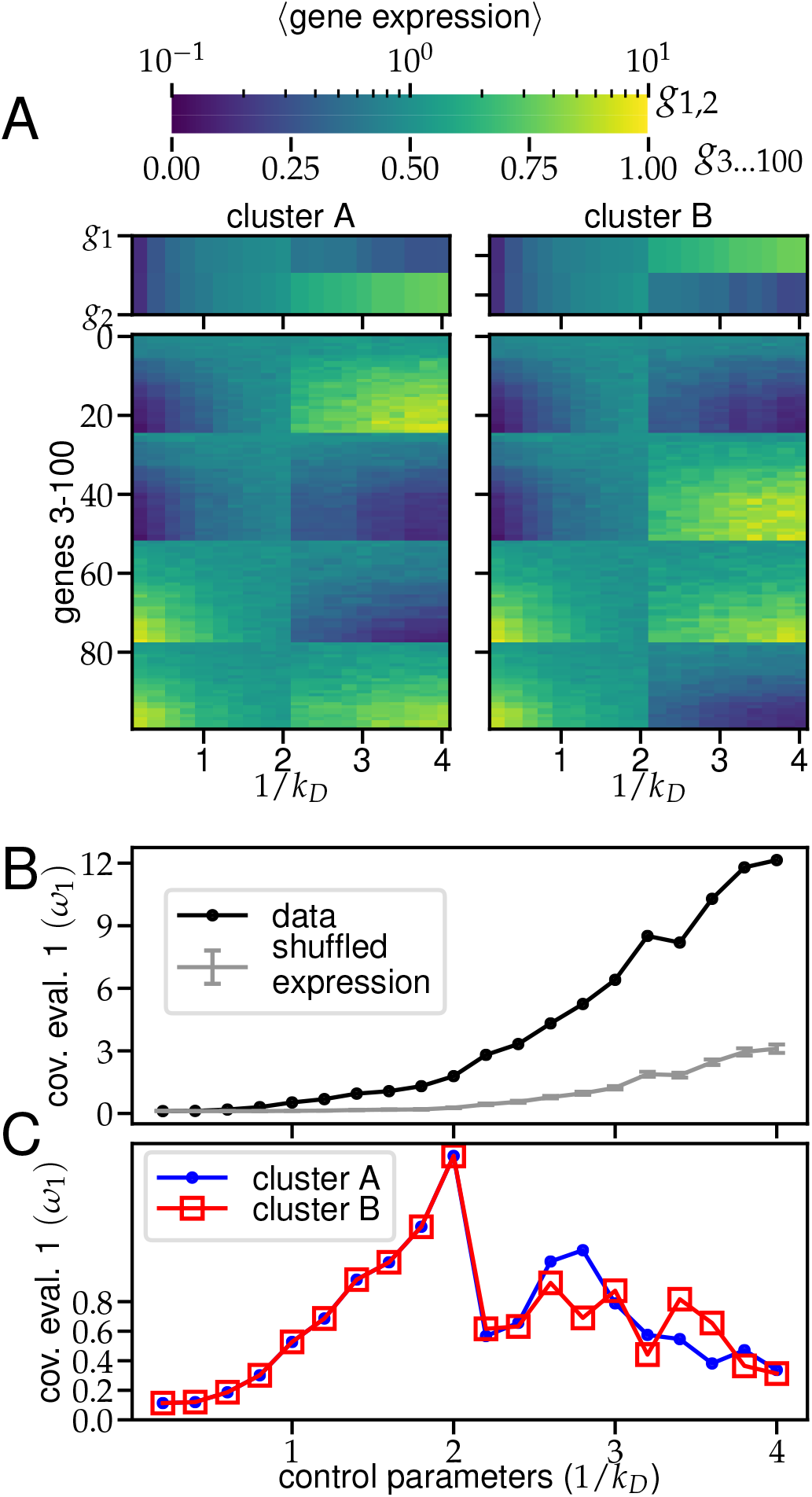
Pitchfork bifurcation analysis. (A) Gene expression as function of the bifurcation variable *τ*, separated by cluster. (B) *ω*_1_ and null for unclustered data. (C) *ω*_1_ for clusters.

**Fig. S5.**
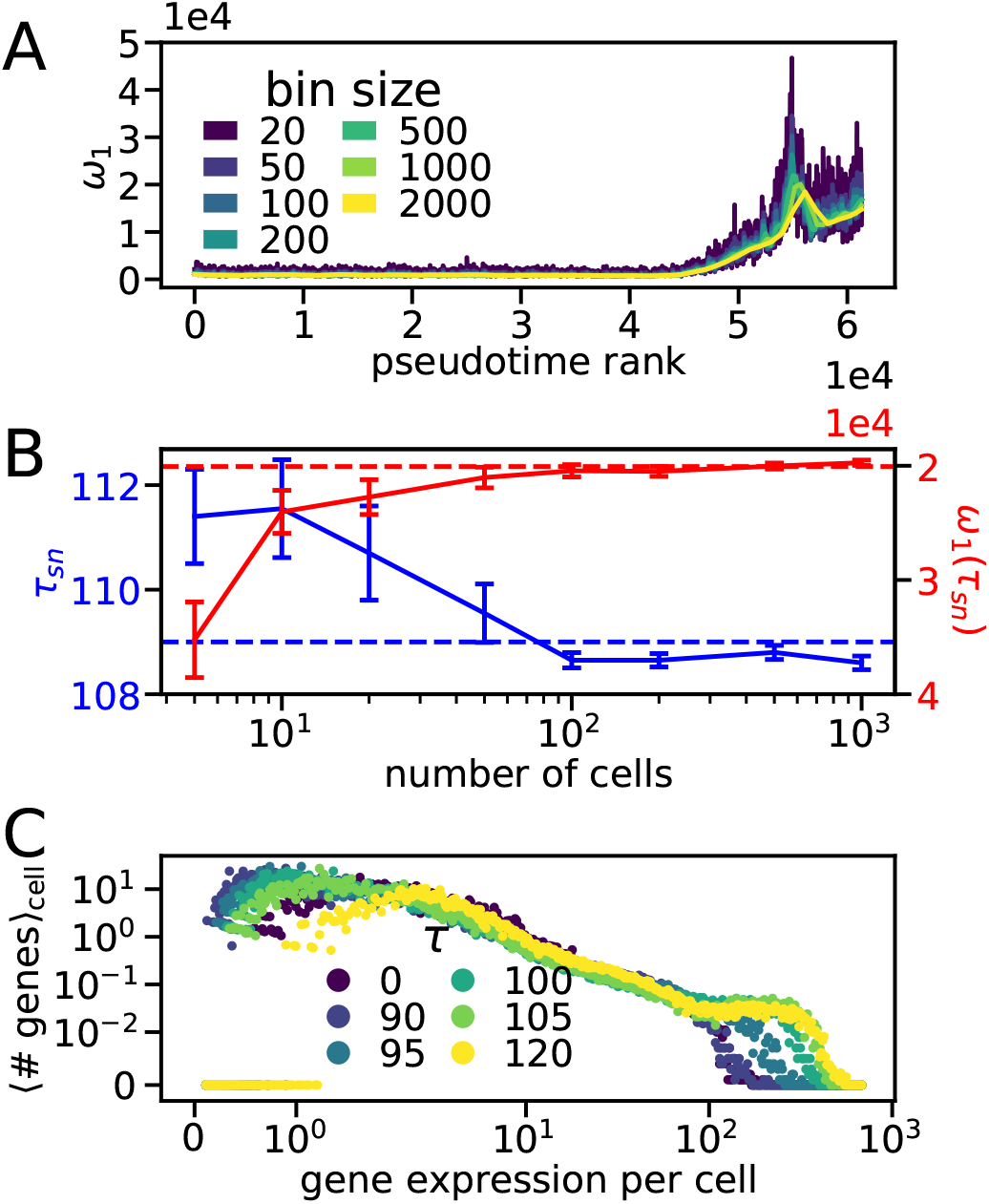
Distributional properties of the neutrophil pseudotime trajectory. (A) Effect of varying the bin size on principal covariance eigenvalue. (B) Effect of undersampling a bin of 1000 cells on the detected saddle-node bifurcation time and magnitude. (C) Distribution of gene expression during the trajectory.

**Fig. S6.**
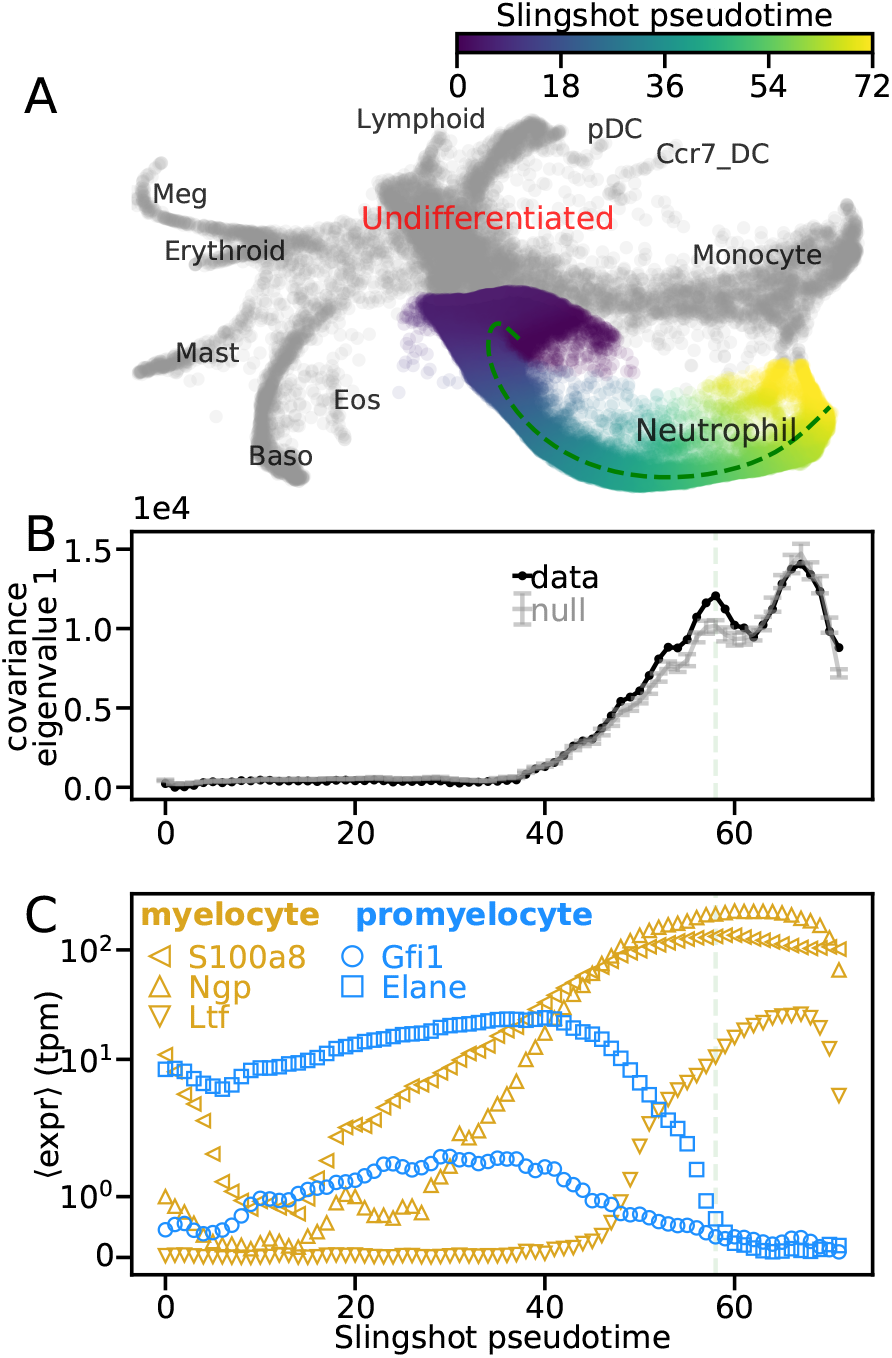
Bifurcation characterization using Slingshot pseudotime algorithm. (A) Neutrophil development obtained by applying Slingshot to hematopoiesis scRNA-seq data (23). Principal curves were approximated to 1000 points by setting approx_points = 1000 in the slingshot function as increasing approx_points further did not affect results. (B) Largest covariance eigenvalue (black) compared with a statistical null (gray, details in Secn. S3) in each 1000 cell pseudotemporal bin, shifted to have 0 min, using the Slingshot pseudotime ordering. Error bars of null are one SD. (C) Average expression of promyelocyte (blue) and myelocyte (gold) marker genes in Slingshot pseudotemporal bins (23). SEM error bars are smaller than symbols. Light green line in (B-C) indicates peak of bifurcation window.

**Fig. S7.**
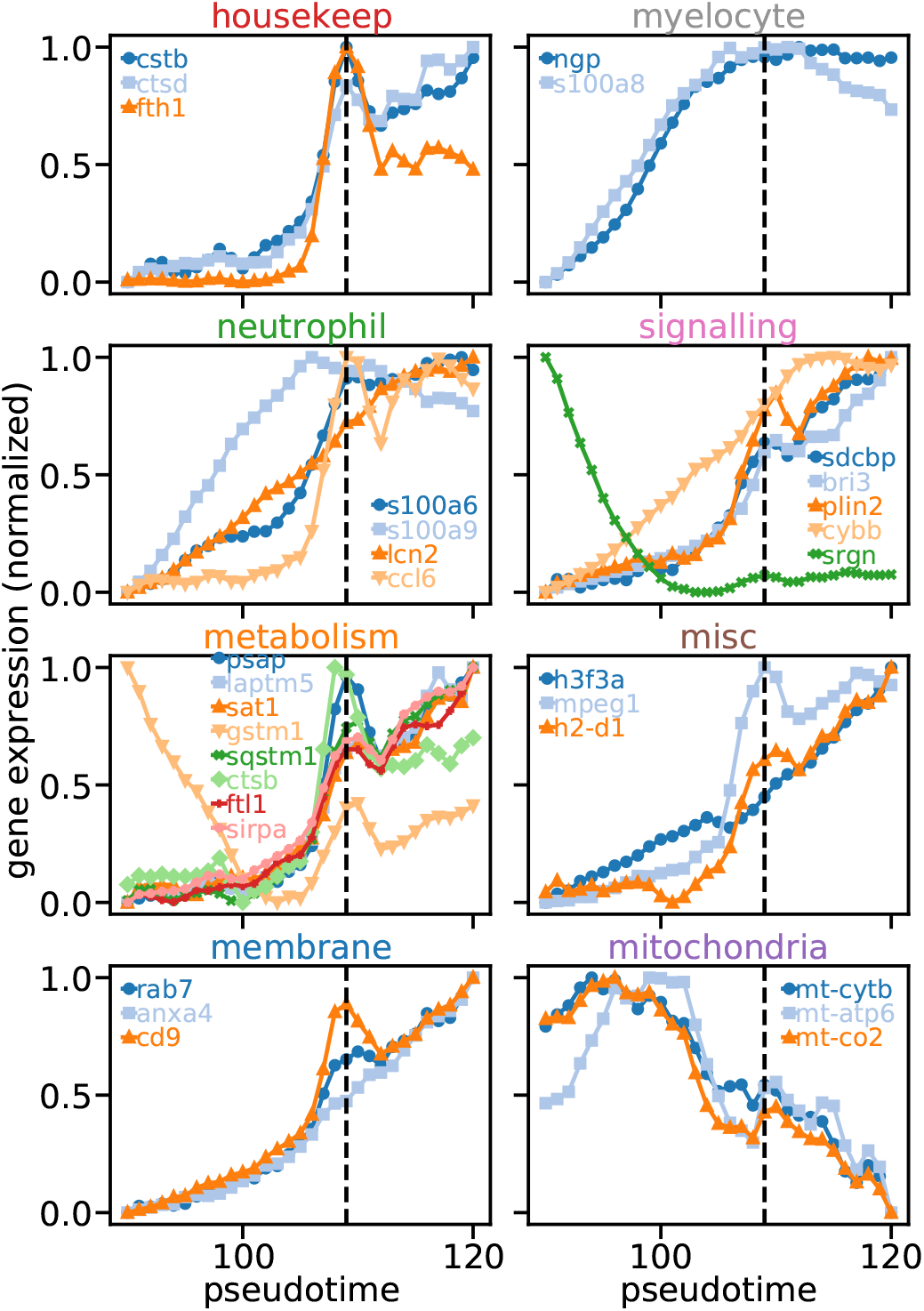
Gene expression of gene groups in Fig. 6, averaged over all cells in bin that have non-zero expression for that gene as a function of pseudotime, min-max normalized in the pseudotime window shown.

**Fig. S8.**
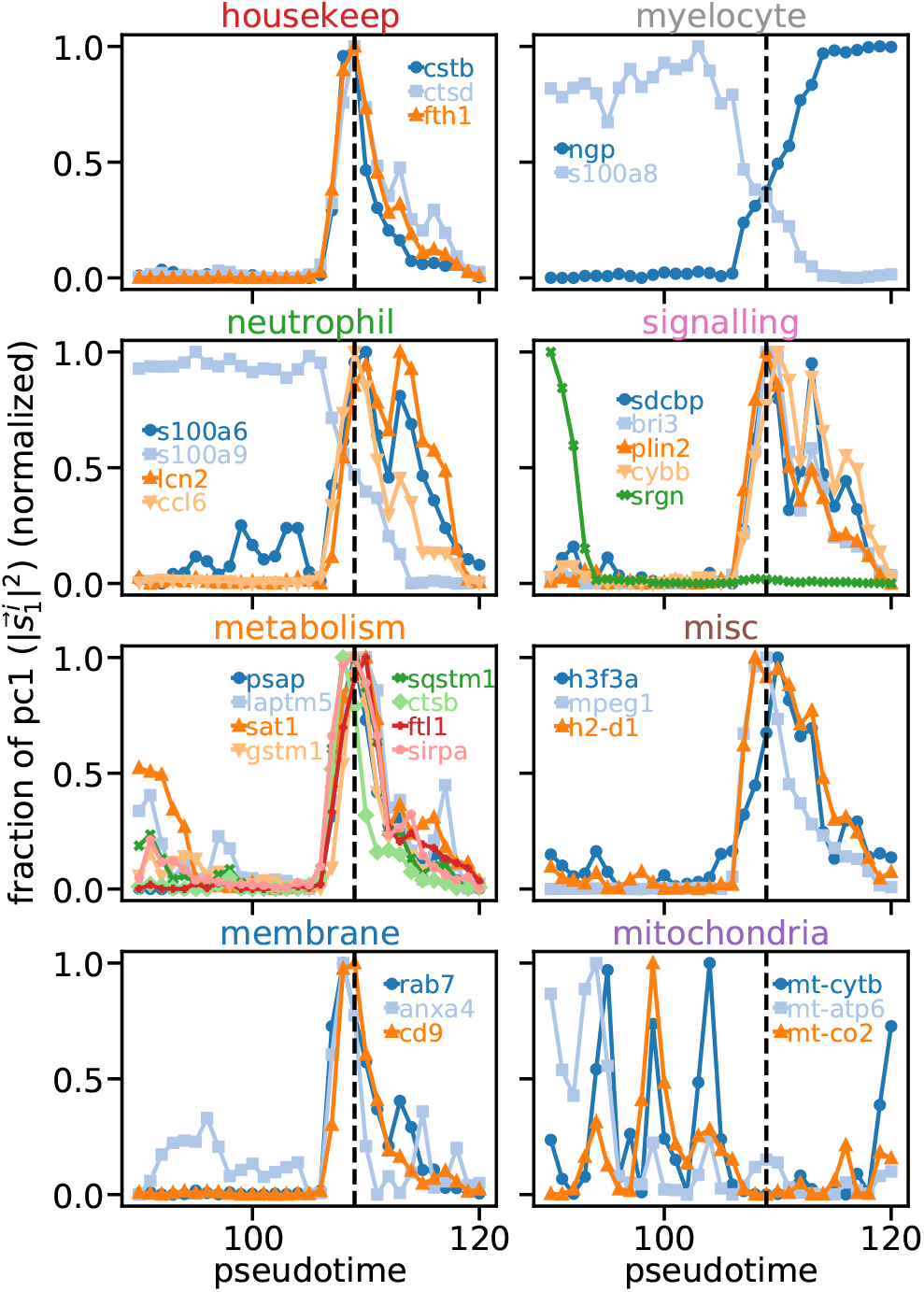
Fractional weight of first covariance eigenvector for gene groups in Fig. 6, as a function of pseudotime, min-max normalized in the pseudotime window shown.

**Fig. S9.**
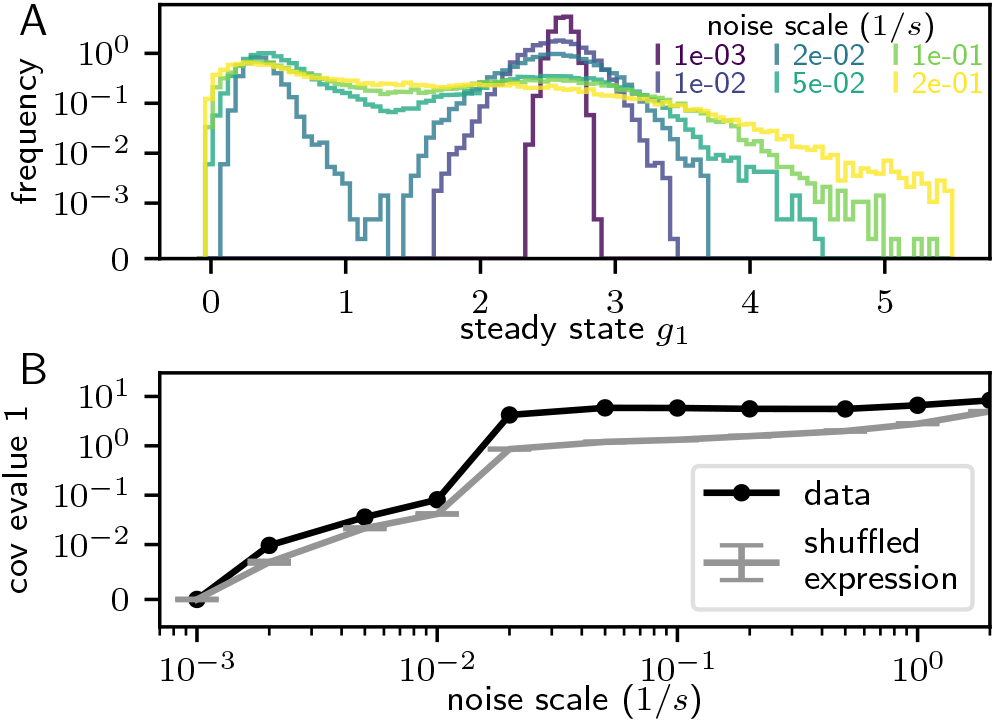
Covariance eigenvalue signature for a noise induced transition. (A) Steady state distributions for the expression of *g*_1_ at varying noise scales. (B) Principal covariance eigenvalue as a function of noise.

